# A biodegradable and restorative peripheral neural interface for the interrogation of neuropathic injuries

**DOI:** 10.1101/2024.08.05.606715

**Authors:** Liu Wang, Tieyuan Zhang, Jiaxin Lei, Shirong Wang, Yanjun Guan, Kuntao Chen, Chaochao Li, Yahao Song, Weining Li, Shimeng Wang, Zhibo Jia, Shengfeng Chen, Jun Bai, Bingbing Yu, Can Yang, Pengcheng Sun, Qingyun Wang, Xing Sheng, Jiang Peng, Yubo Fan, Lizhen Wang, Milin Zhang, Yu Wang, Lan Yin

**Affiliations:** Key Laboratory of Biomechanics and Mechanobiology of Ministry of Education, Beijing Advanced Innovation Center for Biomedical Engineering, School of Biological Science and Medical Engineering, and with the School of Engineering Medicine, Beihang University, Beijing 100191, P. R. China; Institute of Orthopedics, Chinese PLA General Hospital, Beijing Key Lab of Regenerative Medicine in Orthopedics, Key Laboratory of Musculoskeletal Trauma and Injuries PLA, No. 28 Fuxing Road, Beijing 100853, P. R. China; Department of Electronic Engineering, Beijing National Research Center for Information Science and Technology, Institute for Precision Medicine, Center for Flexible Electronics Technology, and IDG/McGovern Institute for Brain Research, Tsinghua University, Beijing, 100084, P. R. China; MegaRobo Technologies Co. ltd, Beijing 100085, P. R. China; School of Materials Science and Engineering, The Key Laboratory of Advanced Materials of Ministry of Education, State Key Laboratory of New Ceramics and Fine Processing, Center for Flexible Electronics Technology, Tsinghua University, Beijing 100084, P. R. China; Department of Dynamics and Control, Beihang University, Beijing 100191, P. R. China

## Abstract

Monitoring the early-stage healing of severe traumatic nerve injuries is essential to gather physiological and pathological information for timely interventions and optimal clinical outcomes. While implantable peripheral nerve interfaces provide direct access to nerve fibers for precise interrogation and modulation, conventional non-degradable designs pose limited utilization in nerve injury rehabilitation. Here, we introduce a biodegradable and restorative neural interface for wireless real-time tracking and recovery of long-gap nerve injuries. Leveraging machine learning techniques, this electronic platform deciphers nerve recovery status and identifies traumatic neuroma formation at the early phase, enabling timely intervention and significantly improved therapeutic outcomes. The biodegradable nature of the device eliminates the need for retrieval procedures, reducing infection risks and secondary tissue damage. This research sheds light on bioresorbable multifunctional peripheral nerve interfaces for probing neuropathic injuries, offering vital information for early diagnosis and therapeutic intervention.

## Introduction

Peripheral nervous systems serve as an important communication network between the central nervous system and the rest of the body, responsible for the transmission of signals for motor output and sensory input ^1,2^. Implantable peripheral nerve interfaces represent an essential device platform that allows direct interrogation of nerve fibers, and can therefore enable enhanced accuracy and selectivity of the monitoring and modulation of neural activities ^1,3^. Potential applications include locomotion restoration ^2,4,5^, pain management ^6^, the treatment of metabolic disorders ^7^, bi-directional prosthetic limb control ^8^, etc.

Peripheral nerve injuries are a prevalent clinical issue, affecting an estimated 13 to 23 individuals per 100,000 annually ^9^. Repairing of severe traumatic nerve injuries continues to be a formidable challenge and the outcomes remain unsatisfactory ^10^. Inadequate treatment can result in permanent motor, sensory, and autonomic impairments. Following nerve transection injuries (e.g., severe injuries or amputation), abnormal nerve tissue regrowth in a disorganized manner often occurs, resulting in traumatic neuroma formation and subsequent chronic pain (as many as 36.5% patients reported chronic neuropathic pain following traumatic nerve injury in the upper extremity) ^11–13^. In cases of amputation, neuroma formation can also greatly complicate prosthetic integration ^12^. This can impact procedures such as targeted muscle reinnervation (TMR), which aims to enhance myoelectric prostheses by transferring transected residual nerves to alternative distant muscles, and disrupted tissue regrowth by neuroma could cause significant difficulties wearing a prosthesis ^14–17^. These challenges arising from unsuccessful reinnervation can impose substantial socioeconomic burdens and psychological distress, greatly impacting patients’ quality of life ^18^.

Since early repair and diagnosis is crucial for optimal therapeutic results ^13^, monitoring the initial stages of healing after severe nerve injuries is essential, which can offer vital physiological and pathological insights for timely interventions and enhanced function restoration ^19–22^. Traditional diagnostic approaches involve physical assessments, imaging technologies (such as ultrasound and computed tomography (CT)), and intraoperative electrodiagnostic tests. However, these methods are limited by challenges related to low spatiotemporal resolution, high cost, frequent clinic visits, and the complexity of real-time monitoring, and therefore hinder early diagnosis ^11,22^. Multifunctional peripheral nerve interfaces present a promising approach for remote and continuous monitoring of nerve injuries, which however remains largely unexplored. By integrating both restorative and monitoring functions, the nerve interface has the potential to promote nerve regeneration and facilitate early detection of recovery challenges during the initial phases of nerve repair. This enables prompt interventions and enhances treatment optimization, ultimately maximizing rehabilitation outcomes and prosthetic integration.

One major challenge in developing neural interface for probing nerve injuries is that conventional peripheral nerve interfaces (e.g., nerve cuffs) are based on non-degradable materials ^23,24^, and often necessitate additional surgeries for subsequent device removal to avoid permanent materials retention, imposing increased infection risks and associated hospital costs. In addition, fibrotic encapsulation typically occurs around the implant especially at the injured site due to foreign body reactions ^25,26^, further complicating the extraction procedure and risking secondary damage to the nerve tissues. Recent developments in bioresorbable materials and devices indicate great potential ^27,28^. Building fully biodegradable and multifunctional peripheral nerve interfaces can offer a promising approach to mitigate the issue by obviating the need for surgical retraction, which however remains unexplored. Such biodegradable electronic platform can enable temporary monitoring and treatment, where early-stage tracking and intervention is crucial to optimize clinical outcomes but becomes unnecessary once regeneration or reinnervation is complete ^29^.

In this paper, we present a biodegradable and restorative neural interface (Bio-Restor) designed for concurrent monitoring and facilitating the healing of long-gap nerve injuries at a finite time frame matched to the critical initial phase of nerve recovery (Figure 1). The device consists of an electrode array for nerve electrical recording and a self-powered galvanic cell that delivers electrical cues to support tissue repair. The interface is built upon a flexible and biodegradable polymer substrate, complemented by a self-morphing shape memory polymer film, which aids the surgical placement of the device onto the injured nerve. The biodegradable neural interface is integrated with a wireless bio-potential recorder for signal amplification, digitization and remote transmission. Employing machine learning techniques, we realize the evaluation of nerve recovery status and the early identification of neuroma formation, and accomplish greatly enhanced therapeutic outcome (Figure 1). A key attribute of this electronic platform is its construction from fully biodegradable materials, which degrades into biologically benign byproducts after the monitoring window, thus eliminating the need for surgical retraction and reducing the risk of tissue damage and infection. The proposed material strategy and device scheme open up new avenues for the interrogation and intervention of nerve injuries to accomplish optimal therapeutic results.

**Figure 1.**
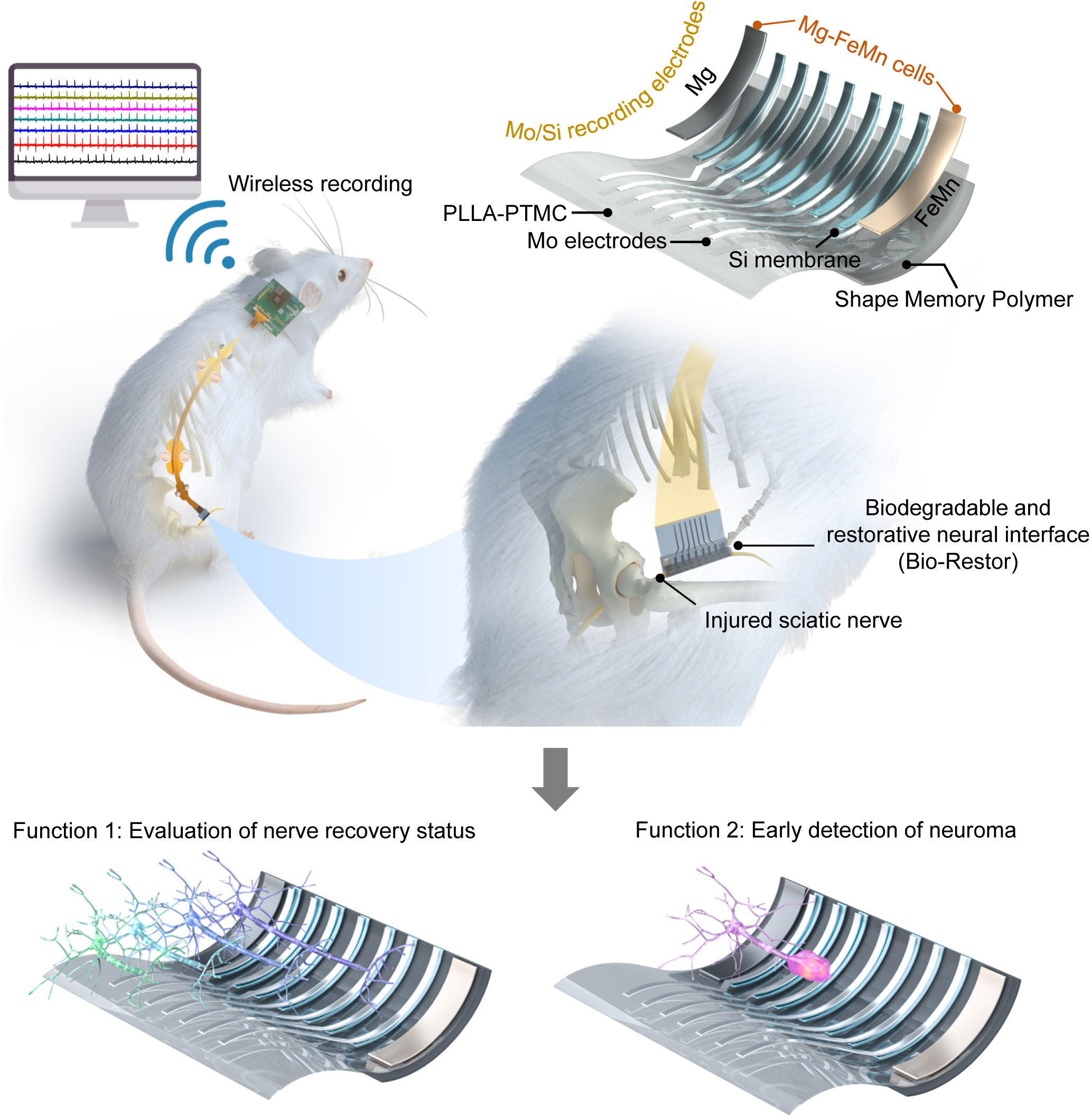
Biodegradable and restorative neural interface (Bio-Restor) for wireless interrogation of neuropathic injuries. Schematic illustration of the device, which is composed of a biodegradable substrate PLLA-PTMC (30 μm, 10 × 10 mm), a Mg-FeMn galvanic cell (Mg, 3.5 μm, 1.5 × 5 mm; FeMn, 1.5 μm, 1.5 × 5 mm), Mo/Si recording electrode array (Mo, 300 nm, 0.3 × 5 mm; Si, 2 μm, 0.5 × 5 mm), and a biodegradable shape memory polymer layer (400 μm, 10 × 5 mm). Evaluation of nerve recover status and early detection of neuroma have been demonstrated based on the biodegradable, restorative and self-morphing neural interface.

## Results

### Materials strategies and device fabrication

The illustration of the device structure of Bio-Restor is depicted in Figure 1, with the corresponding fabrication process given in Figure S1. Specifically, the device includes a 7-channel electrode array for signal recording and a galvanic cell to facilitate functional recovery (Figure 1). The bilayer dissolvable recording electrodes are composed of molybdenum (Mo) thin films (300 nm) and highly doped N-type monocrystalline silicon (Si) membranes (2 µm thick, conductivity 200-1000 S/cm), to ensure desirable conductivity and sufficient operational lifetime. The underlying rationale is based on the slow dissolution rates of highly doped Si, which can serve as effective and conductive water barriers for Mo electrodes ^30,31^. Mo electrodes are deposited through sputtering by a shadow mask on a biodegradable and flexible substrate which is a copolymer of poly(l-lactic acid) and poly(trimethylene carbonate) (PLLA-PTMC, 60:40 ratio, ∼30 μm thick), followed by the transfer-printing of Si membranes to achieve the bilayer electrode array (Figure S1). A biodegradable galvanic cell comprised of thin-film magnesium (Mg, 3.5 µm) and iron-manganese alloy (FeMn, 1.5 µm) electrodes are deposited on the PLLA-PTMC substrate next to the Mo/Si electrode array (Figure 1 and Figure S1), providing electrical cues to promote nerve regrowth, as reported previously^32^. The flexible nature of the neural interface (Figure S2) enables desirable mechanical properties when interfacing with soft peripheral nerves.

An additional biodegradable shape memory polymer (SMP) layer (∼400 μm) consisting of PLLA-PTMC with a different copolymer ratio (70:30) is integrated with the device to facilitate implantation procedure by self-rolling around the peripheral nerve at body temperature (Figure S1). The copolymer ratio of 70:30 is adopted for PLLA-PTMC to ensure that the recovery temperature is adjusted to physiological temperature range ^33^. The device in a rolled-up state fixed by the shape memory polymer appears in Figure 2a, and the recovery from the temporarily flattened state (room temperature, in air) to the final self-curved state (37 °C, in phosphate buffered saline (PBS)) is given in Figure 2b. The differential scanning calorimeter (DSC) measurement of the SMP indicates that the glass transition temperature (T_trans_) is approximately 36.14°C, which confirms the shape memory effects at body temperature. Movie S1 illustrates the deployment of the device utilizing SMP on the intact nerve of Sprague-Dawley (SD) rats by applying PBS at 37 °C.

**Figure 2.**
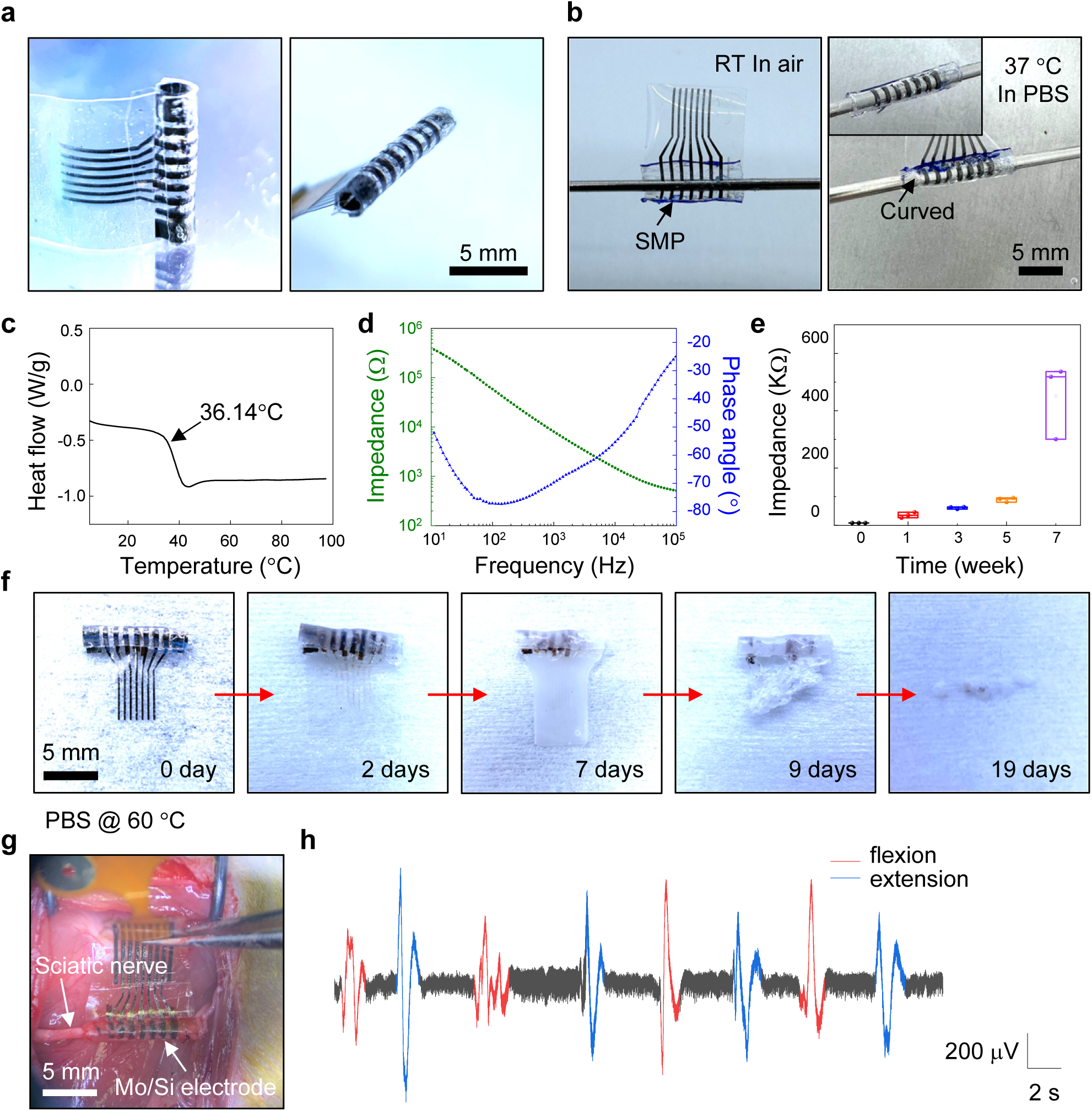
Biodegradable, restorative and self-morphing neural interface. **a**, Image of the biodegradable and restorative neural interface: front view (left) and side view (right). **b**, Neural interface in the temporarily flattened state in air at room temperature (left) and in the permanent rolled-up state after immersion in the PBS (pH 7.4, 37 °C) solution (right). **c**, DSC analysis of the biodegradable shape memory polymer layer. **d**, Representative electrochemical impedance spectra of the biodegradable Mo/Si recording electrode. **e**, Electrochemical impedance at 1 kHz of the biodegradable Mo/Si recording electrode before and after immersion in PBS over 7 weeks. **f.** Images collected at various stages of accelerated dissolution of Bio-Restor in PBS (pH 7.4, 60 °C). **g**, Biodegradable Mo/Si electrode array implanted on the intact sciatic nerve. **h**, Recorded signal traces using the Mo/Si electrode array on sciatic nerves during the flexion-extension movement of the hindlimb. All data are presented as mean ± s.d. of n = 3 rats per condition.

Figure 2d presents the representative electrochemical impedance spectra data for the Mo/Si electrodes in PBS at room temperature with an exposure area of ∼1 mm^2^. The impedance at 1 kHz is 8.1 ± 0.1 kΩ, which is comparable to conventional gold (Au) electrodes with identical dimensions (Figure S3). Moreover, the impedance of Si/Mo electrodes demonstrates reasonable stability after soaking in PBS at 37 °C up to week 5, with an increase in impedance from 35.6 ± 9.8 kΩ (week 1) to 61.1 ± 3.9 kΩ week 3 (Figure 2e). Similar rise in impedance has been reported for non-degradable electrodes ^5,34^. At week 7, a significant increase in impedance is observed, likely resulting from the extensive dissolution of electrode materials. To evaluate the degradation characteristics of Bio-Restor, dissolution is performed in PBS solutions under accelerated conditions (pH 7.4, 60 °C), and the results are given in Figure 2f. The Mg electrodes of the galvanic cell and Mo contacts without Si encapsulation dissolves within 2 days due to relatively rapid degradation. Mo contacts are therefore always coated with biodegradable polymer before implantation to extend the operational time frame. Subsequent disappearance of the FeMn galvanic electrodes and bilayer Mo/Si recording electrodes occurs after approximately 9 days. The polymeric substrate materials degrade through hydrolysis almost entirely after approximately 20 days.

To evaluate the recording performance of the Mo/Si electrodes, we implant the electrode array on the intact sciatic nerves of SD rats (Figure 2g). A commercial neural cuff based on platinum (Pt) electrodes is also investigated as a reference (Figure S4). It is noted that the Mo/Si electrode array can easily wrap around the sciatic nerve due to the incorporation of SMP, enabling a seamless device/tissue interface and minimizing potential tissue damage. Evoked neural signals (compound action potential) is first investigated by applying electrical stimulation at the proximal side (Figure S5). Corresponding results are given in Figure S6 and S7, and the recorded signals are comparable between the cuff electrodes and the biodegradable electrodes (Figure S6). Additionally, reduced stimulation voltage results in reduced signal amplitude, and the evoked neural signals remain similar with inverted stimulation phase (Figure S7). Furthermore, electrical signals recorded on sciatic nerves of SD rats during periodic movement of the hindlimb exhibit a strong correlation with the flexion-extension movement, and representative results are shown in Figure 2h. This is comparable to the signals obtained by the cuff electrodes (Figure S8). Collectively, these results demonstrate that the bioresorbable Mo/Si electrodes are reliable in recording neural signals on sciatic nerves. We then assess the feasibility of utilizing the biodegradable interface for monitoring nerve recovery progress and detection of neuroma to enable timely intervention.

### Biodegradable neural interface for tracking nerve recovery

We integrate a wireless bio-potential recorder with the biodegradable neural interface to enable signal amplification, digitization and wireless data transmission during the free movement of SD rats, as depicted in Figure 3a. Figure 3b presents the specific circuit structure of the wireless circuit, data recording and video capturing system. An analog front end (AFE) with chopper amplification technology is designed to acquire and convert the neural signal to digital data, while maintaining low noise and low power consumption. The design was fabricated with 180 nm technology featuring a die size of 5 mm by 2.95 mm. The measured results demonstrates an integrated input-referred-noise of the low noise amplifier (LNA) of 0.79 μV from 1 Hz to 1 kHz, and a spurious-free dynamic range (SFDR) of 76 dB with a sinusoid input (93 Hz, 14 mVpp), as shown in Figure S9.

**Figure 3.**
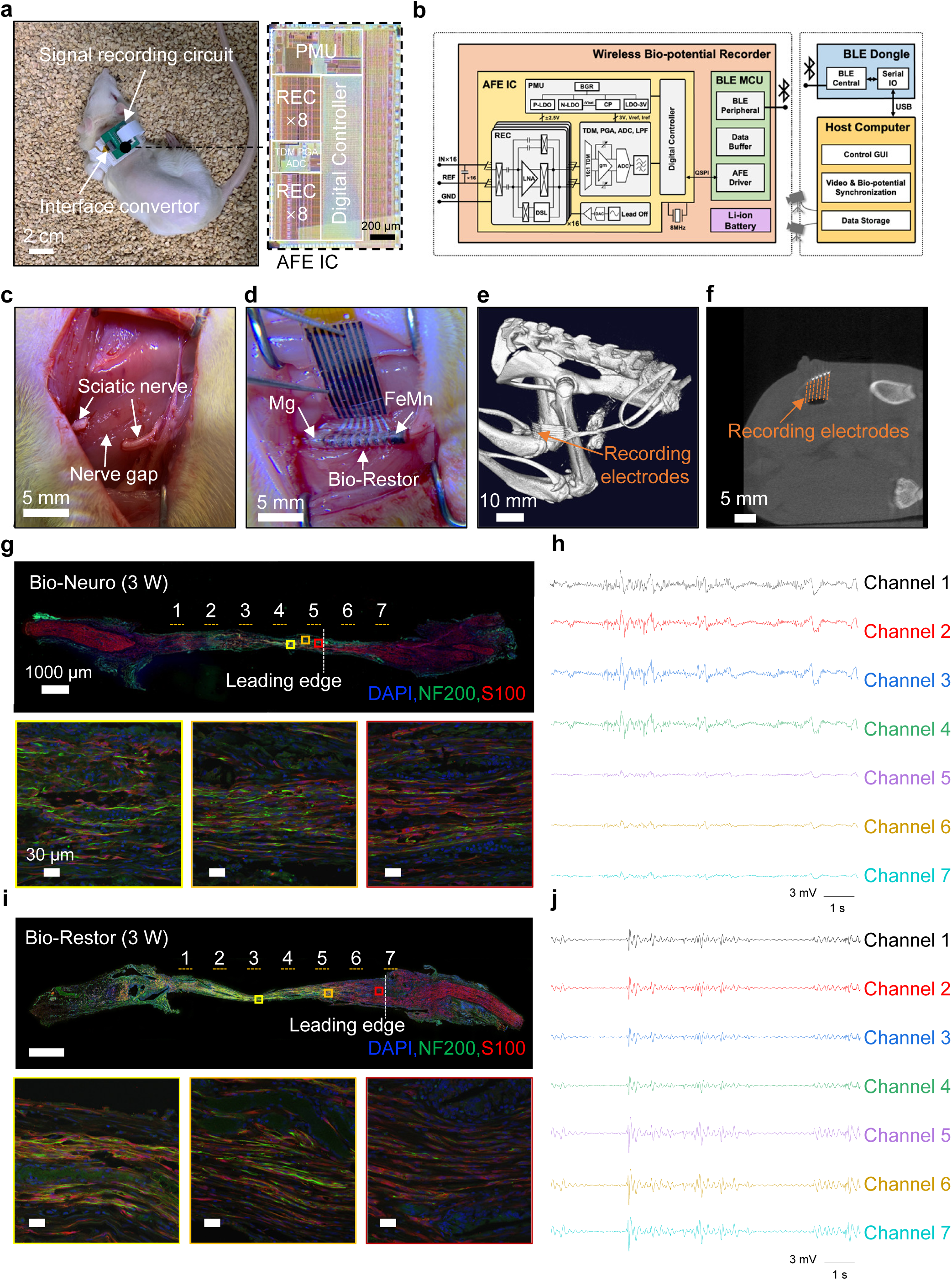
Evaluation of nerve recovery status using Bio-Restor. **a**, Image of a freely moving rat with an implanted Bio-Restor and a wireless module. The enlarged view shows the image of the circuit chip. **b**, Schematic illustration of the wireless circuit, data recording and video capturing system. **c**, Transected sciatic nerve with a 10-mm gap in SD rats. **d**, Bio-Restor implanted at the injured site of nerve gap. **e**, 3D and **f**, cross-sectional micro-computer tomography images of the implanted Bio-Restor. Dotted orange lines indicate the position of connected wires to the multi-channel electrodes. **g**, Immunofluorescent images of the longitudinal section of the nerve segment with implanted Bio-Neuro at 3 weeks postimplantation. Top: immunohistochemical analysis demonstrating the leading edge (white dashed lines) of axonal growth. Yellow dashed lines indicate the position of neural electrodes. Bottom: enlarged view of the immunofluorescent images at 3 different locations. Immunohistochemical staining: axons (NF200, green), Schwann cells (S100, red), and nuclei (DAPI, blue). **h**, Recorded signal traces using Bio-Neuro with rats walking on a treadmill at 3 weeks postimplantation. **i**, Immunofluorescent images of the longitudinal section of the nerve segment with implanted Bio-Restor at 3 weeks postimplantation. Top: Immunohistochemical analysis demonstrating the leading edge (white dashed lines) of axonal outgrowth. Yellow dashed lines indicate the position of neural electrodes. Bottom: enlarged view of the immunofluorescent images at 3 different locations. **j**, Recorded signal traces using Bio-Restor with rats walking on a treadmill at 3 weeks postimplantation. All data are presented as mean ± s.d. of n = 3–4 rats per condition.

To evaluate the capability of tracking nerve recovery in long-gap nerve injuries, we implant Bio-Restor to bridge 10-mm nerve defects in the sciatic nerves of SD rats (Figure S10), with a bipolar configuration to reduce electromyographic noise (Figure S11). The biodegradable neural interface without the galvanic cell is also investigated for comparison, referred to as the Bio-Neuro group (Figure S10). The images of implanted devices and implantation procedure are illustrated in Figure 3c, 3d and Figure S12. The biodegradable neural interface resides at the injured site, routes subcutaneously up to the neck, and connects percutaneously to the wireless circuit for remote signal recording. Rats are able to move freely after device implantation (Movie S2). Figure 3e and 3f displays a micro-computer tomography (micro-CT) constructed 3D image and the cross-sectional view of the implanted device respectively. Wireless signal recording of free-moving rats on a treadmill is conducted at week 2 and week 3 (Figure S13), which represents the critical initial stage of nerve recovery. At week 3, the skin wound of SD rats at the implantation site has fully healed (Figure S14) and the retrieved biodegradable electrodes in the Bio-Restor remain intact (Figure S15). Figure S16 display the images of Hematoxylin and Eosin (H&E) staining of the regenerated axons in the Bio-Restor and Bio-Neuro groups at week 3, indicating no significant inflammation and thus desirable biocompatibility of the device.

Representative recorded signal traces of the Bio-Restor and Bio-Neuro groups and corresponding immunofluorescent images of longitudinal sections of regenerated nerve segments at 3 weeks postimplantation is shown in Figure 3g–j. Interestingly, we can observe a correlation of the leading edge of regenerated axons and the position of recording electrodes that are able to pick up signals. This is because electrodes located distally from regenerated axons are less likely to sense compound neural signals. Specifically, the electrode position is determined by the relative location of the recording electrodes and the footprints of electrodes left on the histological images (e.g., the creation of holes after tissue sectioning in Figure 3g, 3i and Figure S16). The enlarged images in Figure 3g and 3i demonstrate the representative morphology of regenerated axons (green fluorescent area), with the last one depicting the leading edge of axonal growth. In the Bio-Restor group, the front end of axons has reached around the position of electrode 7 (Figure 3i), and all the electrode channels exhibit recorded signals (Figure 3j). By contrast, the control group (Bio-Neuro group) demonstrates the leading edge of regenerated axon has only reached the position around electrode 5 (Figure 3g), and only 4 channels recorded signals with significant amplitude (Figure 3h). The faster restoration of axonal growth in the Bio-Restor group is attributed to the electrical cues offered by the integrated galvanic cell, consistent with previous report^32^. These results suggest that the position of the leading edge of regenerated axons could be qualitatively tracked based on the electrode channels that are able to record significant signals, allowing real-time monitoring of regrowth rates and therefore potentially enabling timely intervention to ensure optimal recovery outcomes.

With the assistance of machine learning methods, we can further analyze the recorded electrical signals during treadmill walking to establish a correlation with gait parameters. These results can provide an important indicator of the recovery status, as nerve injury and regeneration often lead to changes in kinematics and spatiotemporal parameters of gait. Treadmill and Catwalk gait analysis is first conducted using traditional methods, which involve high-speed camera recordings from the lateral and bottom views respectively (Figure 4a). The treadmill gait analysis includes the placement of markers on the hip, knee, ankle, and toe for subsequent DeepLabcut analysis (Figure S17), to obtain essential gait parameters such as joint mechanics, swing phase and stance phase. Specifically, we focus on ankle angle, and the duration of swing phase and stance phase. The stance phase is the period during which the foot is in contact with the ground (Figure S18a), while the swing phase is when the foot raises off the ground (Figure S18b) and swings forward in preparation for the next step. Both the Bio-Restor and Bio-Neuro groups at week 2 and week 3 are investigated, with the gait of normal rats served as the reference. The Bio-Restor group at week 3 exhibits significantly greater ankle angle (p < 0.05) and shorter swing time (p < 0.05) compared to the Bio-Neuro group at week 2. This is evident from the stick diagram in Figure 4b and S19, ankle angle in terminal stance in Figure S20, and actual swing time in Figure 4c. These findings indicate a better recovery in the Bio-Restor group at week 3. Moreover, a comparable pattern is noted in the Catwalk analysis, where the representative footprints of the injured hindlimb in the Bio-Restor group at week 3 appear more intact than those in the Bio-Neuro group at week 2 (Figure 4d and S21). The associated sciatic functional index (SFI), calculated from these footprints, shows a significantly higher value (p < 0.01) in the Bio-Restor group at week 3 compared to the Bio-Neuro group at week 2, signifying improved motor functional recovery.

**Figure 4.**
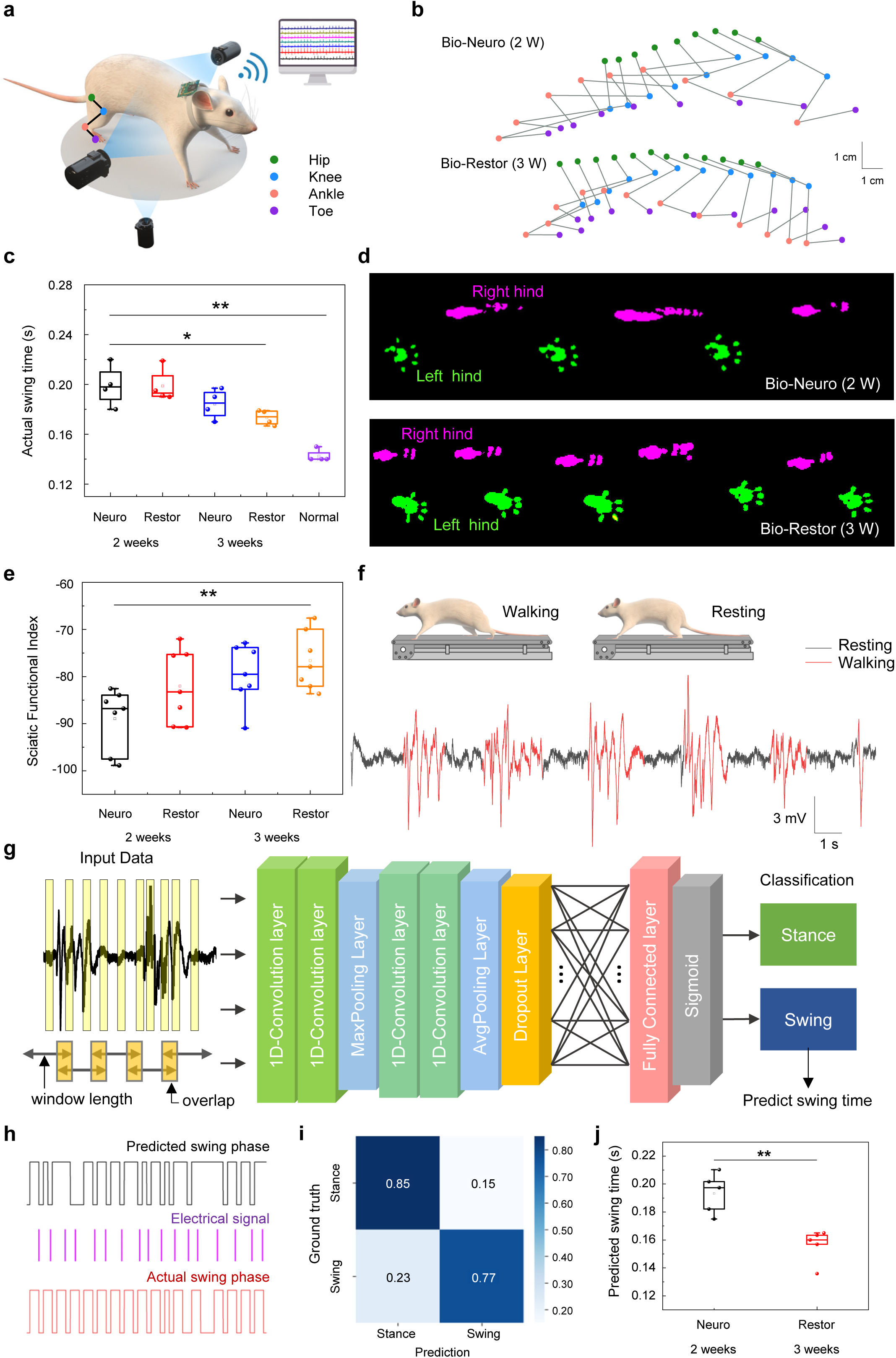
Machine learning assisted evaluation of nerve recovery status through the analysis of swing phase based on recorded electrical signals. **a**, Schematic illustration of conventional gait analysis methods including recording gait from the lateral sides on the treadmill (treadmill gait analysis) and foot prints underneath the runway (Catwalk gait analysis) by high-speed camera. **b**, Kinematic stick diagram decompositions of hindlimb movement on a treadmill. **c**, Statistical analysis of actual swing time of the Bio-Neuro and Bio-Restor groups at 2 and 3 weeks postimplantation. **d**, Representative hindlimb footprints for both injured and contralateral (healthy) sides of the Bio-Neuro group at 2 weeks postimplantation and the Bio-Restor group at 3 weeks postimplantation. **e**, Statistical analysis of sciatic functional index. **f**, Electrical signals recorded by the Bio-Neuro at week 2 during walking and resting states. **g**, Convolutional neural networks (CNN) architecture used to evaluate the swing phase and stance phase based on recorded electrical signals and corresponding gait videos. **h**, Representative results of raster plots of electrical signals, corresponding actual swing phase based on gait videos, and predicted swing phase by CNN architecture of the Bio-Restor group at week 3 postimplantation. **i**, Confusion matrix of the prediction and ground truth of swing and stance phase of the Bio-Restor group at week 3. **j**, Statistical analysis of the predicted swing time of the Bio-Neuro group at week 2 and the Bio-Restor group at week 3. All data are presented as mean ± s.d. of n = 3–5 rats per condition.

Given the notable distinctions in gait results obtained through traditional methods between the Bio-Restor group at week 3 and the Bio-Neuro group at week 2, we proceed to analyze the recorded electrical signal traces of these two groups to establish correlations with gait parameters. This analysis aims to assess the recovery status based on electrical recording. Representative recorded signal traces of the Bio-Neuro group at week 2 are given in Figure 4f and Movie S3, and a correlation is observed between the patterns of electrical signals and the phases of walking or resting. The recorded signal are fed into a Convolutional Neural Network (CNN) to perform swing and stance phase recognition (Figure 4g). Electrical signal dataset was preprocessed before training the network to remove obvious artifacts resulted from movements to improve accuracy. Overlapping windows and oversampling technique are employed for data augmentation and sample balancing. Convolutional layers are applied for feature extracting from the raw acquired neural samples. The pooling and dropout layers are applied to avoid overfitting ^35^. The representative result of the Bio-Restor group at week 3, including predicted swing phase, raster plot based on electrical recording, and actual swing phase obtained by camera recording, is shown in Figure 4h. A reasonable classification accuracy in terms of swing (77%) and stance phase (85%) are achieved (Figure 4i and S22). The classification accuracy of the Bio-Neuro group at week 2 is 72% and 79% for swing and stance phase, respectively (Figure S22). Furthermore, the predicted swing time indicating the recovery progress can be obtained through CNN analysis, and the results of the Bio-Restor group at week 3 and the Bio-Neuro group at week 2 are summarized in Figure 4g. The results indicate a significant reduction of swing time for the Bio-Restor group at week 3 compared to that of the Bio-Neuro group at week 2 (p < 0.01), suggesting a faster walking speed and therefore better recovery. This trend is consistent with the gait analysis obtained by conventional methods (Figure 4b–e). These results altogether suggest that the real-time electrical signals recorded by the bioresorbable interface can be utilized to remotely interrogate the leading edge of axonal growth and gait parameters, therefore predicting the recovery status of injured nerves.

### Biodegradable neural interface for early detection of neuroma

Neuroma often occurs with severe nerve injuries and amputations, and manifests as intricate clusters of disorganized axons encapsulated within fibrous scar tissues at the injured site ^36^. Clinical studies have indicated that neuroma prevalence rates range from 40% to 70% following nerve transection ^37^. The presence of neuromas could lead to subsequent persistence of neuropathic pain and pose formidable obstacles to effective implementation of prosthetic devices in the cases of amputation. Conventional diagnostic methods for neuroma utilize physical assessments, imaging technologies, and intraoperative electrodiagnostic tests. However, the obstacle arises from the challenge of enabling real-time and remote monitoring, thereby impeding early diagnosis. The biodegradable neural interface offers a promising solution to address these issues, allowing early detection of neuroma and timely therapeutic intervention, which is vital for achieving successful reinnervation and optimizing clinical outcomes. To evaluate the feasibility of neuroma detection, we create a 10-mm nerve defects on the sciatic nerves. The Bio-Restor is placed at the proximal side of sciatic nerve for direct signal recording, while liberating the distal nerve stump to allow neuroma formation (Figure 5a). Once neuroma is detected, timely intervention (e.g., surgical excision) is possible, enabling a better recovery (Figure 5a).

**Figure 5.**
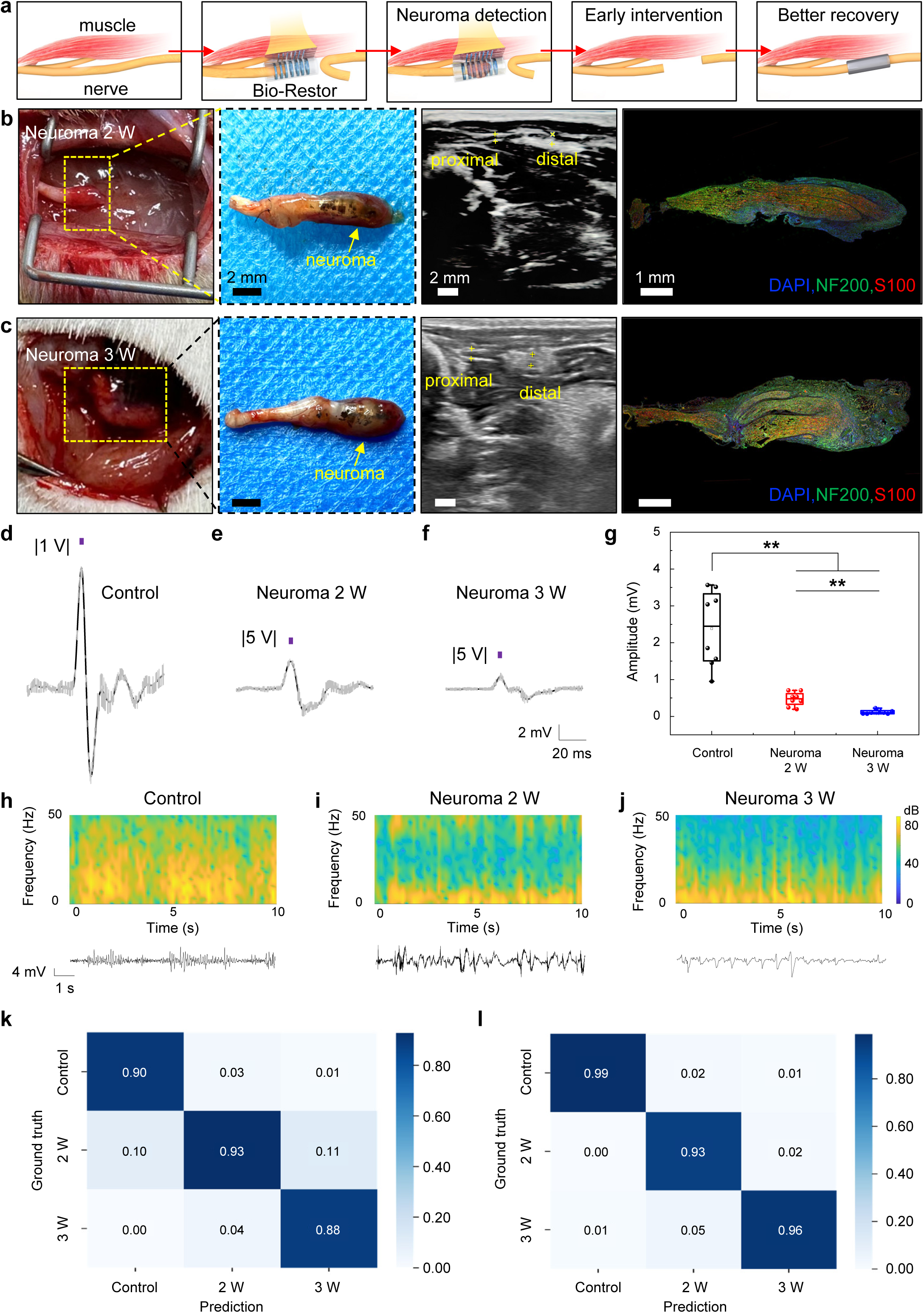
Early detection of neuroma using Bio-Restor. **a**, Schematic illustration of the formation of neuroma, detection by Bio-Restor and early intervention to achieve improved therapeutic outcomes. **b**, Photographs, ultrasound and immunofluorescent images of neuroma at 2 weeks post injuries. **c**, Photographs, ultrasound and immunofluorescent images of neuroma at 3 weeks post injuries. **d–f**, Representative recorded evoked signals by Bio-Restor of the control (intact nerve), neuroma at 2 weeks, and neuroma at 3 weeks respectively, under the electrical stimulation at the proximal side with a voltage amplitude of 1 V, 5 V and 5 V respectively. **g**, Statistical analysis of the amplitude of evoked neural signals. **h–j**, Representative recorded electrical signals and associated time-frequency spectrograms of rats walking on a treadmill of the control (regenerated nerves at 3 weeks), neuroma at 2 weeks, and 3 weeks respectively. **k**, Confusion matrix of the prediction and ground truth of the control, neuroma 2 W and neuroma 3 W groups based on evoked neural signals. **l**, Confusion matrix of the prediction and ground truth of the control, neuroma 2 W and neuroma 3 W groups based on electrical signals recorded during walking on a treadmill. All data are presented as mean ± s.d. of n = 3 rats per condition.

At 2 weeks post nerve injuries, the formation of enlarged and bulbous nerve tissues are observed (optical images in Figure 5b), indicating the occurrence of neuroma. Immunofluorescent staining of the nerve tissues reveals chaotic and disorganized axonal growth (displayed as the green fluorescent area in Figure 5b) instead of the typical directional neurites seen in healthy nerves, further confirming the formation of neuroma. This group is referred as the Neuroma 2 W group. An increase in the size of neuromas is observed at 3 weeks post nerve injuries (Figure 5c), referred as the Neuroma 3 W group. Notably, identification of the occurrence of neuroma at such an early stage cannot be unambiguously determined with traditional ultrasound techniques (ultrasound images in Figure 5b, c), and tissue harvesting is needed for morphological and immunohistochemical analysis to confirm its presence. In contrast, the biodegradable nerve interface is able to identify significantly different electrical signal patterns at week 2 and week 3, achieving early neuroma detection. Firstly, we investigate evoked neural signals in the presence of neuroma, and representative results are given in Figure 5d–f. We study signal traces recorded by the Bio-Restor device in the control (intact nerve), neuroma 2 W and neuroma 3 W groups, upon electrical stimulation on the proximal site. The results show a significant reduction in amplitude with the presence of a neuroma, and an increase in neuroma size leads to a further decrease in amplitude (Figure 5g). This is likely due to the fibrotic encapsulation of disorganized neural tissues, which greatly blocks electrical signals. Clinical studies have reported that poor recovery of neuroma-in-continuity is associated with significant decrease in the amplitude of nerve action potential during intraoperative measurement ^38^, which further validates the evoked neural recording in our work. Due to the sensitivity of electrical signals to passivation, these findings indicate that the neural interface implanted around the injured nerve can provide a robust tool for identifying neuroma formation. In comparison to the traditional electrophysiological assessment performed on target muscles, which necessitates extensive time for reinnervation to elicit a visible response, directly examining electrical activity at the injury site allows for much earlier detection ^38^.

Furthermore, electrical signal traces recorded during treadmill walking are investigated. With the biodegradable neural interface, real-time monitoring can be accomplished remotely without the need of additional devices (e.g., stimulation electrodes). Representative results and associated time-frequency spectrograms are given in 5h–j, for the control (regenerated nerves without neuroma formation at week 3), neuroma 2 W, and neuroma 3 W groups. The results indicate a significant decrease in the relative energy within the high-frequency band (approximately 15-50 Hz) associated with the formation of the neuroma (neuroma 2 W and neuroma 3 W groups). This is likely caused by the presence of thick fibrotic tissues due to neuroma formation, similar to the mechanism that causes the decrease in the amplitude of evoked neural signals. The distinct patterns of electrical signals can be analyzed to enable early diagnosis of neuroma. With the assistance of machine learning analysis (CNN), a high-accuracy of classification can be accomplished and the results are given in Figure 5k and 5l. The identification accuracy of neuroma formation at week 2 is 93%, based on signals recorded during evoked stimulation and treadmill walking respectively. In comparison, traditional ultrasound technique is unable to detect neuroma formation at such an early stage (Figure 5b). Moreover, the identification accuracy for Neuroma 3 W is 88% and 96%, respectively, based on signals recorded during evoked stimulation and treadmill walking. These results suggest that the biodegradable interface can provide essential information for early diagnosis of neuroma formation.

Early detection of neuroma is crucial to enable timely intervention and enhance therapeutic efficacy. To demonstrate the importance of early diagnosis, we investigate the outcomes of intervention at different time points. Specifically, upon the detection of neuroma at 2 weeks post nerve injuries, we follow a standard clinical treatment, by performing excision of the neuroma and employing a nerve conduit to guide nerve regrowth (intervention at 2 W) (Figure 6a). For comparison, the same intervention at a later stage, 5 weeks post nerve injuries (intervention at 5 W), is performed. The outcomes of gastrocnemius muscle atrophy and nerve conduction recovery at 12 weeks post-intervention are evaluated. Figure 6b presents the representative image of gastrocnemius muscle of both the injured and contralateral (healthy) sides, suggesting significant muscle atrophy with a later intervention (5 W). To quantitatively assess muscle atrophy, the muscle wet weight ratio is calculated by comparing the gastrocnemius muscles on the injured side with those on the contralateral healthy side, as shown in figure 6c. The results indicate that the muscle wet weight ratio of the intervention at week 2 is significantly greater than that of the intervention at week 5 (p < 0.05). Figure 6d displays the Masson trichrome staining of gastrocnemius muscles in the injured limbs, revealing notable reduction in collagenous deposits and larger average cross-sectional muscle fiber area in the early intervention (2 W) group. Statistical analysis of the cross-sectional area of muscle fibers demonstrates that the muscle fiber area in the early intervention group (2 W) are significantly greater than that of the later intervention group (5 W) (p < 0.01) (Figure 6e). Moreover, nerve conduction recovery is examined by assessing compound muscle action potential (CMAP), which is recorded on the targeted gastrocnemius muscles upon the electrical stimulation of the proximal sciatic nerve stumps on the injured side. The representative results are illustrated in Figure 6f. Statistical analysis reveals a significant improvement in CMAP amplitude in the early intervention group (2 W) compared to the later intervention group (5 W) (p <0.05). In all, these results suggest the essential benefits of early neuroma detection and intervention, which can greatly facilitate nerve reinnervation and reduce muscle atrophy.

**Figure 6.**
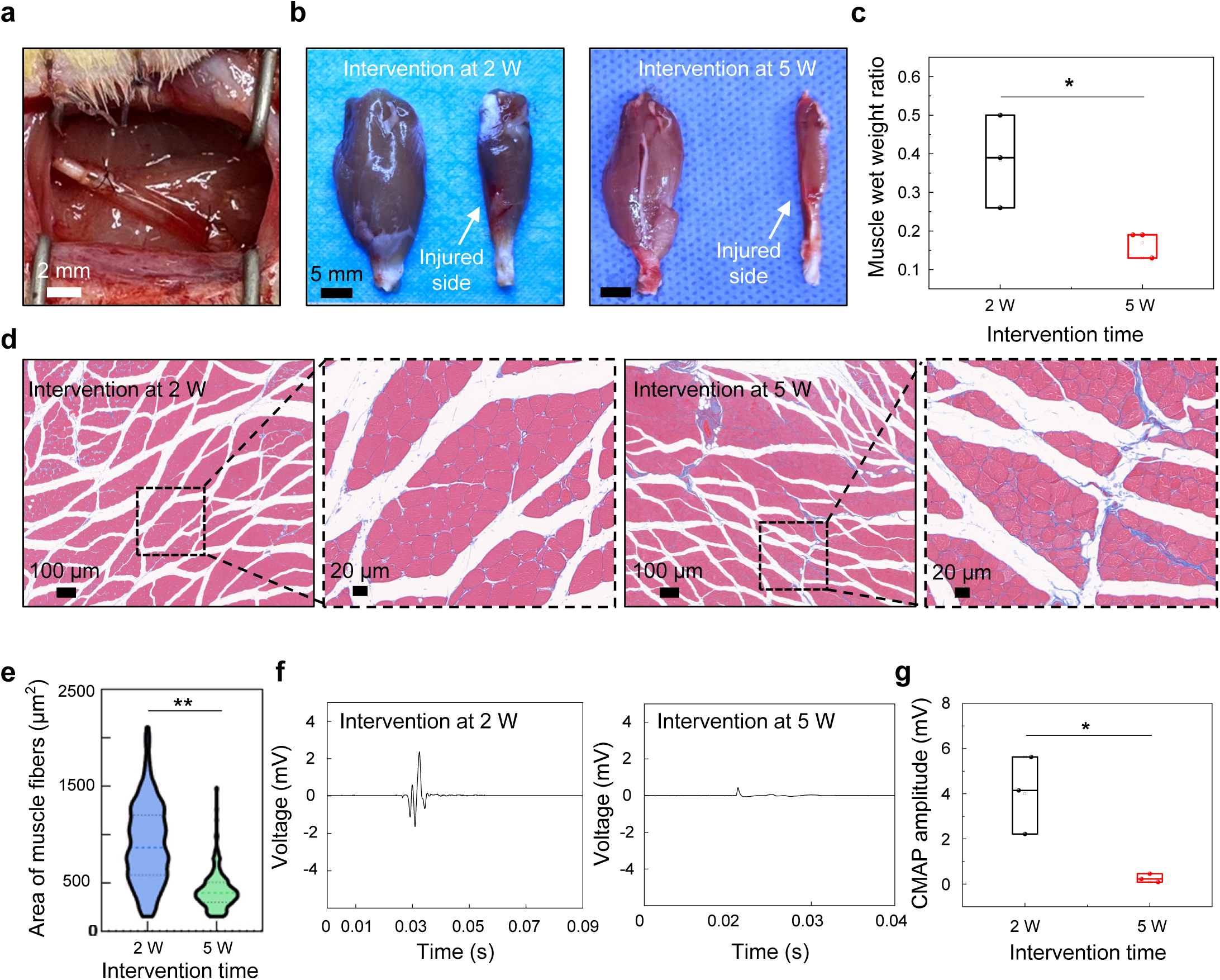
Therapeutic outcomes of early intervention of neuroma 12 weeks postoperatively. **a**, Photograph of the treatment of neuroma by excision and nerve guidance conduits. **b**, Photographs of the gastrocnemius muscles of the contralateral (healthy, left) and injured side (right) with intervention at week 2 and week 5. **c**, Statistical analysis of the wet weight ratio of the gastrocnemius muscles of the injured hindlimb. **d**, Masson’s trichrome staining images of the transverse sections of muscles of the injured hindlimb. **e**, Statistical analysis of the cross-sectional area of muscle fibers of the injured hindlimb. **f**, Representative CMAP at the injured side. Electrical stimulation (1.0 mA, 1 Hz, 0.1 ms) is applied at the proximal nerve stumps. **g**, Statistical analysis of CMAP amplitude at the injured side. All data are presented as mean ± s.d. of n = 3 rats per condition.

### Biodegradation of the neural interface

The essential feature of the proposed neural interface is that it is constructed with all biodegradable materials and can therefore be resorbed by the body over time after tracking nerve recovery, eliminating surgical complications for device retrieval and minimizing secondary damages to delicate nerve tissues. The active components including Mo/Si electrodes and the galvanic cell largely degrade after 7 weeks of implantation, and the remaining polymeric substrates degrades over a much longer time frame through hydrolysis (Figure 7a). The morphology of the retrieved device, isolated from regenerated nerve tissues, is displayed in the inset of Figure 7a, illustrating a gradual degradation process. The recorded electrical signals remain stable at week 2 and week 3, which provides a sufficient window for monitoring the critical stage of long-gap nerve injuries. The signal starts to decay at week 5 and completely lost the recording capability at week 7 (Figure 7b). Moreover, biochemical analyses of the blood samples of SD rats with device implantation indicate the absence of physiological abnormalities 3 weeks postimplantation with a reference to the control group (Figure 7c). Histological examinations of vital organs, including the heart, liver, spleen, lung, and kidney tissues, reveal no significant adverse effects compared to the control group, suggesting desirable biocompatibility of the biodegradable neural interface (Figure 7d).

**Figure 7.**
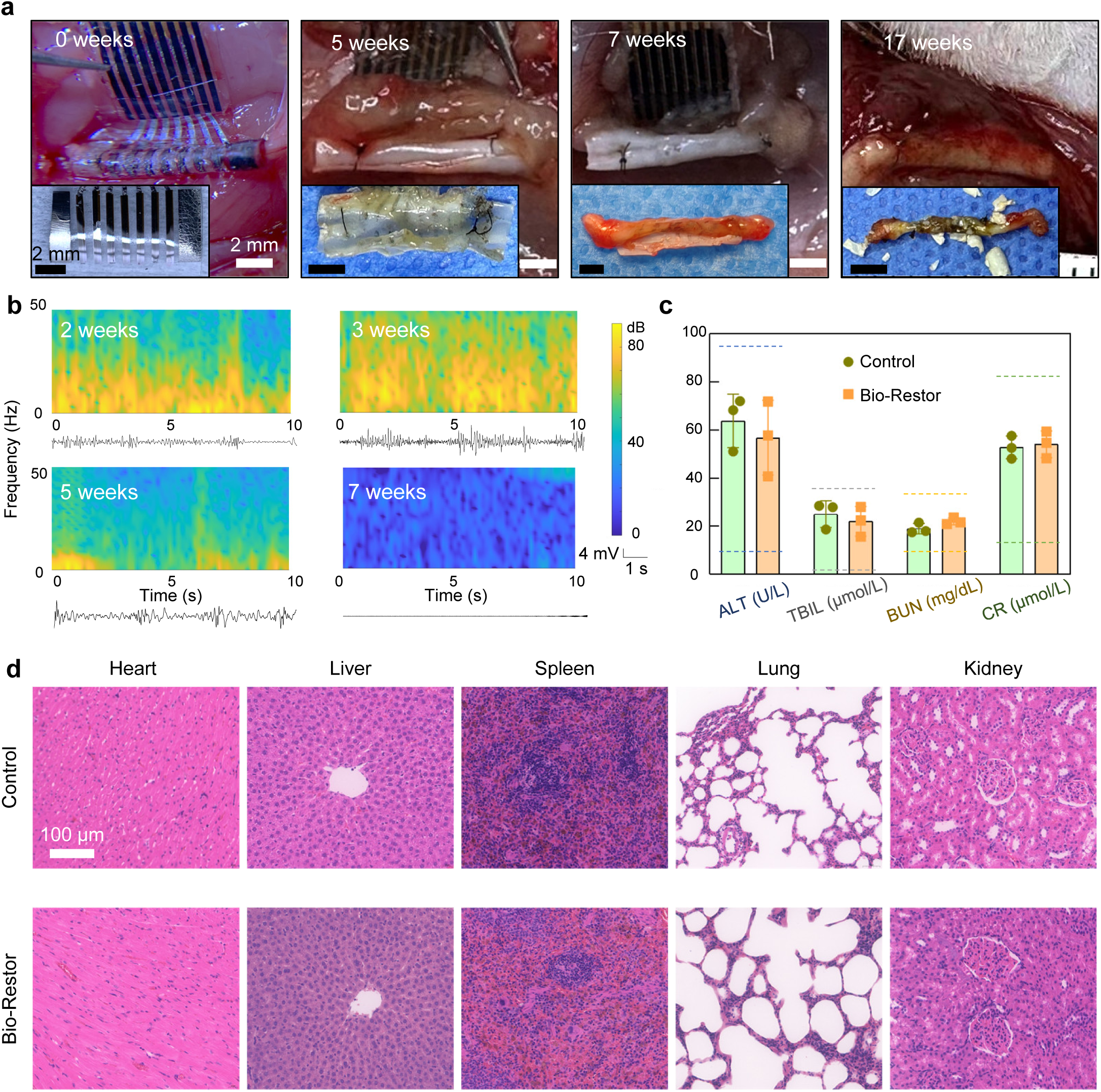
Biodegradable and biocompatible characteristics of Bio-Restor. **a**, Images of the Bio-Restor implanted at the injured sites of sciatic nerves in rodents at different stages of biodegradation over a time frame of 17 weeks. Inset: retrieved device isolated from regenerated nerve tissues. **b**, Representative recorded signal traces and corresponding time-frequency spectrograms at different stages during the course of biodegradation. **c**, Analysis of blood chemistry and blood counts of rats of the control (no device implantation) and Bio-Restor groups at 3 weeks postimplantation. ALT (Aminotransferase); TBIL (Total Bilirubin); BUN (Blood Urea Nitrogen); CR (Creatinine). Dash lines represent the normal range. **d**, H&E-stained histological sections of heart, liver, spleen, lung, and kidney of the control and Bio-Restor groups at 3 weeks postimplantation. All data are presented as mean ± s.d. of n = 3 rats per condition.

## Discussions

We introduce a biodegradable and restorative peripheral neural interface that not only supports nerve regrowth but also enables real-time electrical recording at the critical early stage following neuropathic injuries. We demonstrate the capability of remote monitoring nerve recovery status by tracking the leading edge of axonal growth and the prediction of gait parameters. Furthermore, early detection of the formation of traumatic neuroma is accomplished, which allows timely intervention to significantly improve therapeutic outcomes. The neural interface is constructed with bioresorbable materials that can degrade into harmless byproducts after the monitoring period, negating the need for surgical removal and reducing the risk of complications. The proposed materials strategies and device schemes provide new avenues to interrogate previously unexplored electrical neural signals for real-time monitoring and identification of abnormities following nerve injuries, which could greatly improve the efficacy of treating long segment nerve injuries. The device platform also has potential applications for amputees to facilitate TMR procedures, which can assist reinnervation, identify potential neuroma formation and therefore facilitate earlier and more effective prosthetic integration. Future research direction includes increasing the channel count to enhance the precision and selectivity of signal recording, materials options to achieve controllable operational time frame, and the incorporation of neural modulation component to achieve closed-loop system for enhanced recovery and pain management. Collectively, the research offers new insights into examining electrical activity in the context of post-traumatic nerve injuries or neural disorders, which can uncover crucial information that can optimize treatment protocols, ultimately leading to maximized clinical outcomes and enhanced patient care.

## Materials and methods

### Fabrication of biodegradable and restorative neural interface

Polymer films (7:3 PLLA-PTMC of SMP: 440 μm, 10 × 5 mm and 6:4 PLLA-PTMC: 30 μm, 10 × 10 mm) were fabricated by dissolving PLLA-PTMC (Jinan Daigang Biomaterial Co. Ltd., China) into trichloromethane (CHCl_3_) (Beijing Tongguang Chemical Co. Ltd., China) with a weight-to-volume (w/v) ratio of 1:10. The solutions were then drop-casted and cured for 12 hours at 4 °C to prevent the formation of bubbles. The SMP was then applied onto a needle (diameter, 1 mm) and subjected to heat treatment in an oven at 80 °C for a duration of 30 minutes. This process allowed the polymer to be shaped into a tubular structure. Subsequently, we reopened the tubular structure to a planar form at room temperature, and laminate it with the PLLA-PTMC (6:4) substrate using a minimal amount of CHCl_3_ solution.

Mo recording electrode array (300 nm, 0.3 × 5 mm) were deposited on the PLLA-PTMC layer material through a patterned shadow mask in a magnetron sputter (Beijing Zhongjingkeyi Co. Ltd., China) with a deposition speed of 1.5 Å s^-1^ (370 V, 0.3 A). Mg thin-film anodes (3.5 μm, 1.5 × 5 mm) and FeMn alloy cathodes (1.5 μm, 1.5 × 5 mm) were deposited on two sides of the bilayer material using magnetron sputter system through patterned shadow masks at a deposition speed of 1.8 Å s^-1^ (280 V, 0.23 A) and 1.1 Å s^-1^ (340 V, 0.25 A), respectively. The FeMn alloy is deposited by a customized target with 30 wt % Mn. Highly doped monocrystalline Si membranes (2 μm) were obtained using a silicon on insulator (SOI) wafer (top Si layer: conductivity 0.1–2×10^4^ S·m^-1^, 2 μm). The top Si layer was patterned using a lithographic process followed by reactive ion etching with sulfur hexafluoride gas (SF_6_). Subsequently, Si membranes on top of the SOI wafer were released by hydrofluoric acid (49 % HF, ACS grade, Aladdin). Polydimethylsiloxane (Dow Corning Sylgard 184 kit, 1:10 weight ratio) stamps were applied to transfer the freestanding Si membranes onto the PLLA-PTMC substrate with the Mo electrode array to achieve Mo/Si bilayer recording electrodes.

### Characterization of shape memory performance, electrochemical properties, and biodegradability

Differential scanning calorimetry (DSC) (Naichi Co. Ltd., Germany) was employed to determine the thermal properties of SMP. Nitrogen gas was used as a purging gas at a flow rate of 20 mL/min. The sample underwent an initial heating process, where it was heated from 0 to 100 °C at a rate of 10 °C/min and held at 100 °C for 2 minutes. Subsequently, it was cooled down to 0 °C at a cooling rate of 10 °C/min. This first heating process aimed to eliminate any thermal history effects arising from different materials. Following this, a second heating process was conducted to generate the DSC curve. The glass transition temperature (Tg) was determined from the second heating run.

The electrochemical impedance of the Mo/Si bilayer recording electrodes were investigated using a Gamry Interface 1000E Potentiostat. The measurements were conducted at room temperature in PBS. A three-electrode configuration was employed, with the Si/Mo recording electrodes serving as the working electrode, Ag/AgCl as the reference electrode, and Pt as the counter electrode. The electrochemical impedance spectroscopy (EIS) data were acquired by sweeping the frequency from 10 Hz to 0.1 MHz, which is relevant for neural sensing. A sinusoidal AC excitation signal with an amplitude of 5 mV and a bias of 0 V relative to the open-circuit voltage was applied for the measurement.

To evaluate degradation, the device was immersed in PBS (replaced every day) at 60°C in a water bath. Time-series images were captured at various stages using an optical microscope.

### Signal recording under evoked conditions

To evaluate the performance of neural recording, the biodegradable Mo/Si electrode array and commercial Pt cuff electrodes (Kedou (Suzhou) Brain-computer Technology Co. Ltd, China) were investigated on intact sciatic nerves of SD rats (n = 3 per condition). All animal study procedures followed the institutional guidelines of the Chinese PLA General Hospital (approval number: 2016-x9-07). All trials were performed with adult female SD rats (220-250 g). Surgery was performed under 1.5% isoflurane (RWD Life Science Co., Shenzhen, China) anesthesia. Under sterile conditions, the right hindlimb was dissected to expose the sciatic nerve. To deploy the biodegradable Mo/Si electrode array to the sciatic nerve, a small amount of PBS at 37 °C was used to induce self-rolling of the SMP, ensuring a conformal contact with the nerve tissues. Evoked neural signals were recorded upon the application of electrical stimulation (frequency:1 Hz, amplitude: ± 1 and 0.25 V) at the proximal side of the sciatic nerve. Signal traces were also recorded during the flexion-extension movement of the hindlimb.

### Surgical procedure of device implantation

Long-gap sciatic nerve injuries: Twenty Female SD rats (220-250 g) were anesthetized using 3% pentobarbital sodium solutions (30 mg per kg body weight). The right sciatic nerve was exposed and the surrounding connective tissue was isolated. A 10-mm sciatic nerve defect was created proximal to the branch of the sciatic nerve using a microsurgery scissor. Bio-Restor or Bio-Neuro was connected to the nerve defect area and sutured using 8-0 polypropylene sutures (n = 5 per condition). The receiver routes subcutaneously along the spine to the neck. Finally, muscle layers and the skin were closed with 4-0 silk sutures.

Creation of traumatic neuroma: Eight female SD rats (220-250 g) were anesthetized using 3% pentobarbital sodium solution (30 mg per kg body weight). The right sciatic nerve was exposed, and the surrounding connective tissue was isolated. The nerve was cut 5 mm proximal to the sciatic nerve branch. The proximal nerve stump was enveloped with Bio-Restor and secured to the nearby muscle with 10-0 sutures. The distal nerve stump was folded back and sutured beneath the muscle layer with 10-0 sutures. The wound was then closed in successive layers. The formation of neuroma was examined at 2 weeks and 3 weeks postimplantation by optical observation, ultrasound, immunofluorescent staining (n = 4 per condition).

Intervention of traumatic neuroma: At 2 or 5 weeks following the primary surgery to create the neuroma, a secondary surgery was performed to repair the neuroma on the right sciatic nerve (n = 4 per condition). The base around the proximal nerve was meticulously dissected to examine the enlarged neuroma formation. The distal nerve was isolated from the muscle. The proximal neuroma and the scared area of the distal nerve were excised to reveal the fresh nerve stumps and create a 10 mm nerve gap. 8-0 sutures were utilized to connect the PLLA-PTMC conduit to the nerve defect region. Finally, the wound was closed layer by layer.

### Gait analysis

Treadmill gait analysis: A customized treadmill setup and high-speed camera from the lateral sides was utilized to evaluate the walking patterns of rodents of the Bio-Restor and Bio-Neuro groups at week 2 and week 3 postimplantation (n = 5 per condition). Gait analysis was performed on normal rats for comparison. Markers were placed on the hip, knee, ankle, and toe of the rats as reference points for training of the algorithm. The gait parameters including joint angle, swing time and stance time were extracted from the acquired video stream. A deep neural network (DeepLabcut), consisting of a pretrained ResNet-50 backbone for feature extracting and deconvolutional layers for upsampling ^39^, is employed to identify the locations of the reference points, with less than 5 pixels average error. Joint angles can be calculated from the coordinates of the reference points, and the swing and stance time can be measured by the toe height curve.

CatWalk gait analysis: CatWalk gait analysis was performed for the Bio-Restor and Bio-Neuro groups at week 2 and week 3 after surgery (n = 5 per condition). Briefly, walking tracks were recorded by a high-speed camera located beneath the runway using the CatWalk XT 10.6 gait analysis system (Noldus Information Technology, Wageningen, The Netherlands). Information about the foot contact area, walking posture, and 3D plantar pressure distribution was collected at the same time. At least three identifiable runs were performed for each rat in each measurement. Finally, SFI was calculated using Noldus Catwalk Analysis software.

### In vivo data recording

In vivo signals were recorded for SD rats walking on a treadmill (n = 3 per condition). Evoked neural signals from the neuroma were recorded upon the application of electrical stimulation (duration: 50 μs, frequency: single pulse, amplitude: 5 V) to the proximal side of the sciatic nerve (n = 3 per condition).

### Wireless circuit for signal recording and data transmission

The wireless circuit (20 × 20 × 4 mm) is based on a customized 16-channel analog-front-end (AFE) chip with flexible channel configurations. The nRF5340 Micro Control Unit (MCU) from Nordic Semiconductor is selected to drive the AFE and implement the BLE 5 protocol. The BLE 5 protocol with Data Length Extension (DLE) enables the real-time transmission of the collected electrical data. The entire wireless circuit consumes ∼50 mW of power, which can be supplied by a 100 mAh battery (20 × 20 × 5 mm) with a lifespan of 9 hours. A 120-fps camera enables global shutter, preventing the potential image distortion when the experimental subjects are moving fast.

### Data analysis of recorded signals

Signal processing: A Butterworth high-pass filter at 10 Hz is used to remove low-frequency motion artifact noise, while a low-pass filter with a 300 Hz cut-off frequency removes high-frequency environmental noise. An IIR notch filter addresses industrial frequency noise and its harmonics at 50 Hz and its multiples (100 Hz, 150 Hz, 200 Hz). Wavelet denoising is also applied for signal processing and noise reduction.

Data decoding for gait prediction: In order to decode the neural signals of treadmill walking to establish a correlation with gait parameters, a 1D-CNN model is proposed to classify the acquired neural signals into two classes, including swing phase and stance phase. Based on gait patterns obtained by high-speed camera, electrical signal dataset was preprocessed before training the networks to remove the artifacts resulted from movements and muscle activities to improve accuracy. Overlapping windows and oversampling technique were employed for data augmentation and sample balancing. We adjusted the length of window and the overlapping ratio to test the performance of the classifier. The electrical signal dataset is divided into three subsets, 80% data for training, 10% data for validation and 10% data for testing after preprocessing and normalization. The swing phase and stance phase can be identified from the electrical signal segment using 1D-CNN networks. Subsequently, the swing time and stance time can be calculated. The raster plot is achieved by drawing vertical lines at the corresponding time point to represent neural firing events.

Data decoding for neuroma identification: In order to decode the neural signals as well as to analyze neuroma growth, a 1D-CNN model was applied. The neural signal dataset of control, neuroma (2 W) and neuroma (3 W) was preprocessed before training the network to remove potential artifacts to improve the signal-to-noise ratio using the aforementioned methods. Neural signals collected in two different scenarios, including electrical stimulation and treadmill walking, were used for training and testing the network to comprehensively demonstrate the reliability of the results. After preprocessing and normalization, the electrical signals were segmented into non-overlapping 2-second segments, with 80% used as the training set, 10% as the validation set, and 10% as the test set. The 1D-CNN model can identify neural signal segments as either without neuroma (control), neuroma (2 W), or neuroma (3 W).

### Histological and Immunohistochemical evaluation

At 2 and 3 weeks postimplantation, rats were euthanized with carbon dioxide gas (n = 4 per condition). Sciatic nerve tissue and devices were explanted and fixed in 4% paraformaldehyde for 48 hours. A frozen slicer (Leica, Wetzlar, Germany) was used to cut nerve tissues into 10-μm sections longitudinally. H&E and immunofluorescence staining were performed respectively. The following primary and second antibodies were used: Mouse anti-Neurofilament 200 antibody (N0142, Sigma), Rabbit anti-S100 antibody (ab52642, Abcam), Goat Anti-Mouse IgG H&L (Alexa Fluor® 488, ab150117, Abcam) and Goat Anti-Rabbit IgG H&L (Alexa Fluor® 594, ab150084; Abcam). Finally, the stained sections were scanned using a slide scanning system (3DHISTECH, Budapest, Hungary). All assessments were conducted in triplicate for each animal samples.

### Evaluation of gastrocnemius muscles

Three SD rats from each group were chosen for assessment at 12 weeks post-surgery to analyze their electrophysiological response, wet weight ratio, and muscle fiber area in the gastrocnemius muscle. Motor nerve function was evaluated using a multi-channel physiological signal recording and processing system (RM6240EC, Chengyi). Under anesthesia (1% sodium phenobarbital), the sciatic nerves on the injured side were exposed. Electrical stimulation (3.0 mA, 1 Hz) was administered to the proximal nerve stumps, and CMAPs were recorded from the gastrocnemius muscle to assess motor nerve function. The peak amplitudes of CMAPs were measured and compared between the different groups. Following the recordings, the gastrocnemius muscles from both the injured and unharmed hindlimbs were excised and weighed to determine the wet weight ratio. The muscles were then fixed (4% paraformaldehyde, 12 hours), processed into thin paraffin sections (5 μm in thickness), and subjected to Masson staining for analysis. The stained sections were examined using a slide scanning system (3DHISTECH, Budapest, Hungary), and the cross-sectional area of the gastrocnemius fibers was quantified using Image-Pro Plus 6.0 software.

### Micro-CT and ultrasonography

Micro CT of anesthetized rats was performed on a CT system (Lenovo Co. Ltd., China). Images were acquired with 70 kV, and the projection data were reconstructed with the Workstation. Ultrasound examination was performed at 2 and 3 weeks after the neuroma creation surgery. Rats were anesthetized with isoflurane, and the right hindlimb was shaved to fully expose the skin. High-frequency ultrasound equipment with a 21-to 44-MHz linear array probe (MX4000, Vevo 3100, Visualsonics, Canada) imaged the proximal stump of the sciatic nerve and measured the diameter of the enlarged nerve tissues.

### In vivo biodegradation studies

For in vivo biodegradability evaluation, Bio-Restor was implanted in rats, which were subsequently exposed at 5, 7, and 17 weeks post-implantation. The corresponding Bio-Restor samples were then extracted, and photographs were taken to study the biodegradation behavior in vivo.

### Assessment of hematology and blood chemistry

At 3 weeks postimplantation, rats were deeply anaesthetized and peripheral blood was collected from the orbital vein using vacuum blood collection tubes containing procoagulant and separator gel (QX0028, Servicebio). The blood samples were kept at room temperature for 2 hours, and then the serum was separated in a centrifuge at 3000 rpm for 15 minutes at 4 °C. Biochemical and electrolyte tests were performed by Servicebio Biotechnology Co. LTD.

### Statistical analysis

Statistical analysis was conducted using GraphPad Prism (version 6.0) for two-group comparisons with an unpaired t-test, and SPSS software (version 23.0) for multi-group comparisons using one-way ANOVA. Significance levels were set at *P < 0.05 and **P < 0.01. Data are presented as mean ± standard deviation.

## Acknowledgements

The project was supported by the National Natural Science Foundation of China (T2122010 and 52171239 to L.Y., 32101088 to L.W., U20A20390, 52272277 to X.S. and 11827803 to Y.F., 11932003 to Q.W.), Beijing Municipal Natural Science Foundation (Z220015 to L.Y., Z240017 to L. W.), Beijing Nova Program and Fundamental Research Funds for the Central Universities.

## Author contributions

L.W. and L.Y. conceived the idea. L.W. and K.C. performed material design, fabrication and characterization. J.L., Y.S., W.L., S.W. and M.Z. performed wireless circuit design, fabrication and characterization. T.Z., Y.G., C.L., Z.J., S.C., J.B. and L.Y. designed and performed biological experiments. S.W., L.W., Q.W., B.Y., C.Y., P.S. and L.Y. participated data analysis discussion. L.W., Y.F., J.P., Y.W., X.S. and L.Y. provided tools and supervised the research. L.W., T.Z., J.L., Y.S., W.L., M.Z., X.S. and L.Y. wrote the paper in consultation with the rest of the authors.

## Competing interests

The authors declare no competing interests.

---

**Figure S1.**
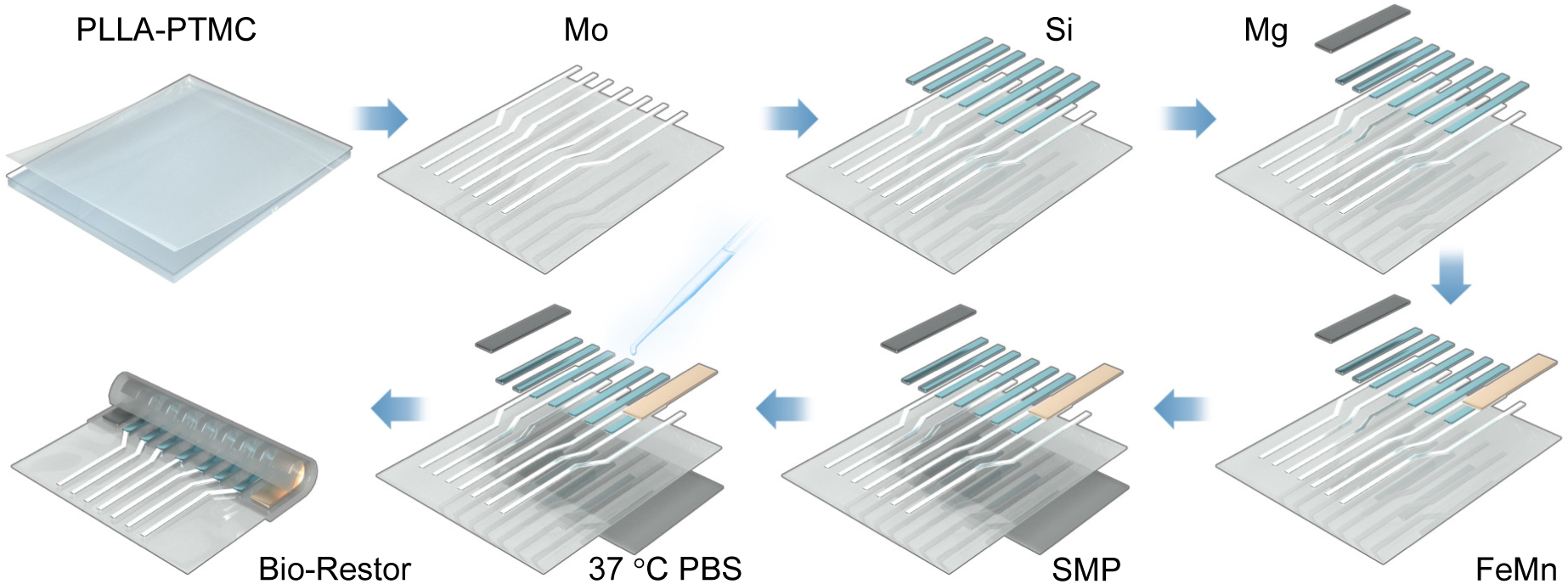
Schematic illustration of processing flow for the fabrication of Bio-Restor.

**Figure S2.**
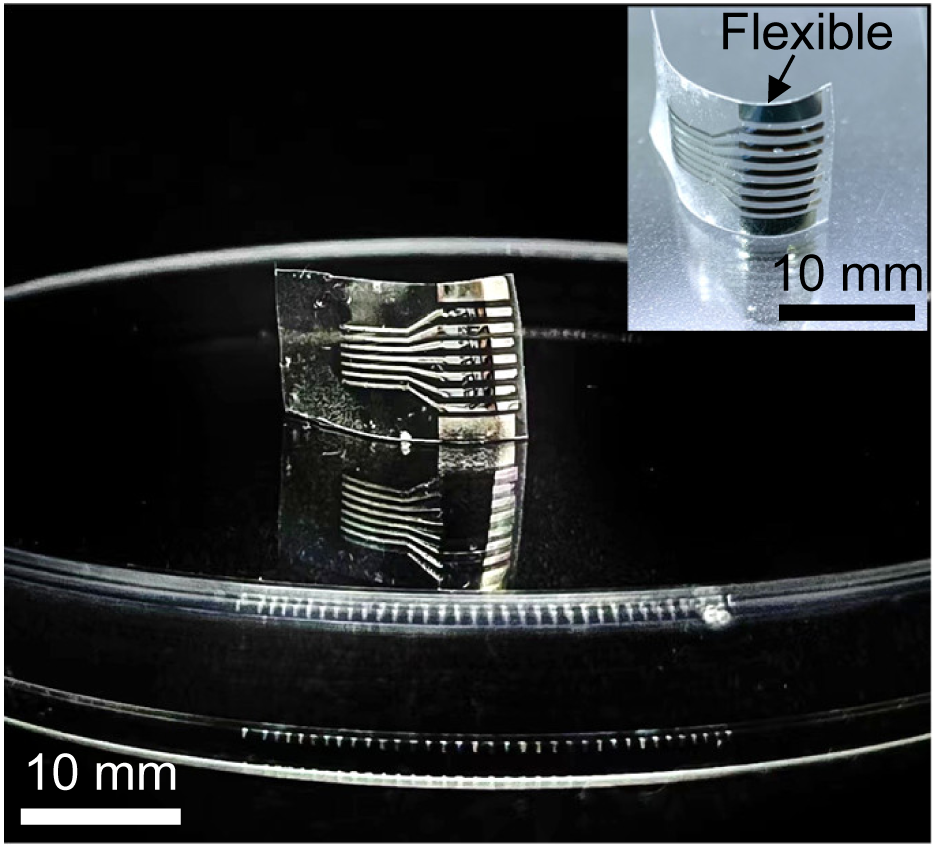
Flexible Bio-Restor in the planar format.

**Figure S3.**
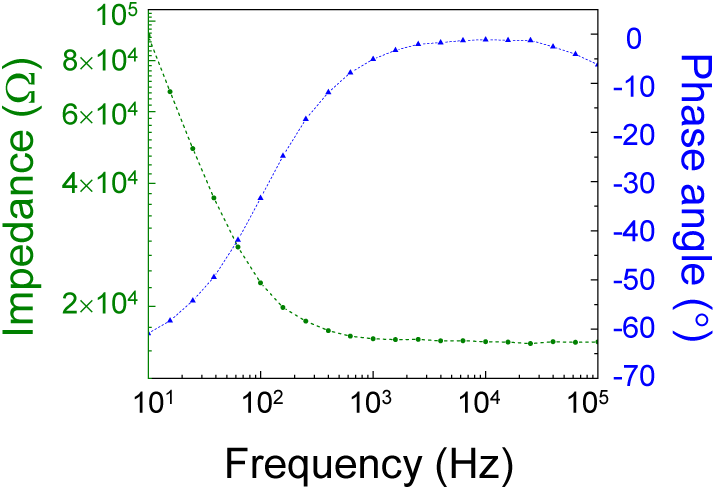
Representative electrochemical impedance spectrum of Au recording electrode.

**Figure S4.**
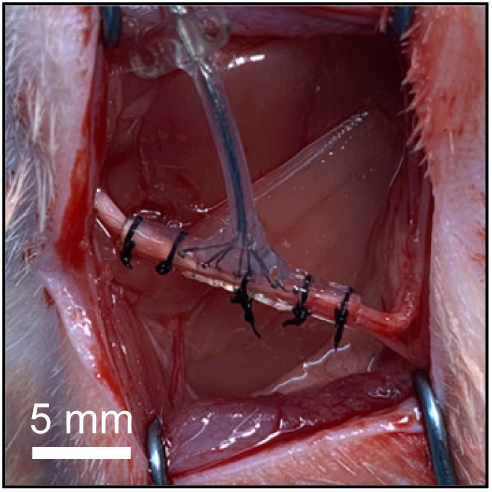
A commercial cuff electrode implanted on the intact sciatic nerve.

**Figure S5.**
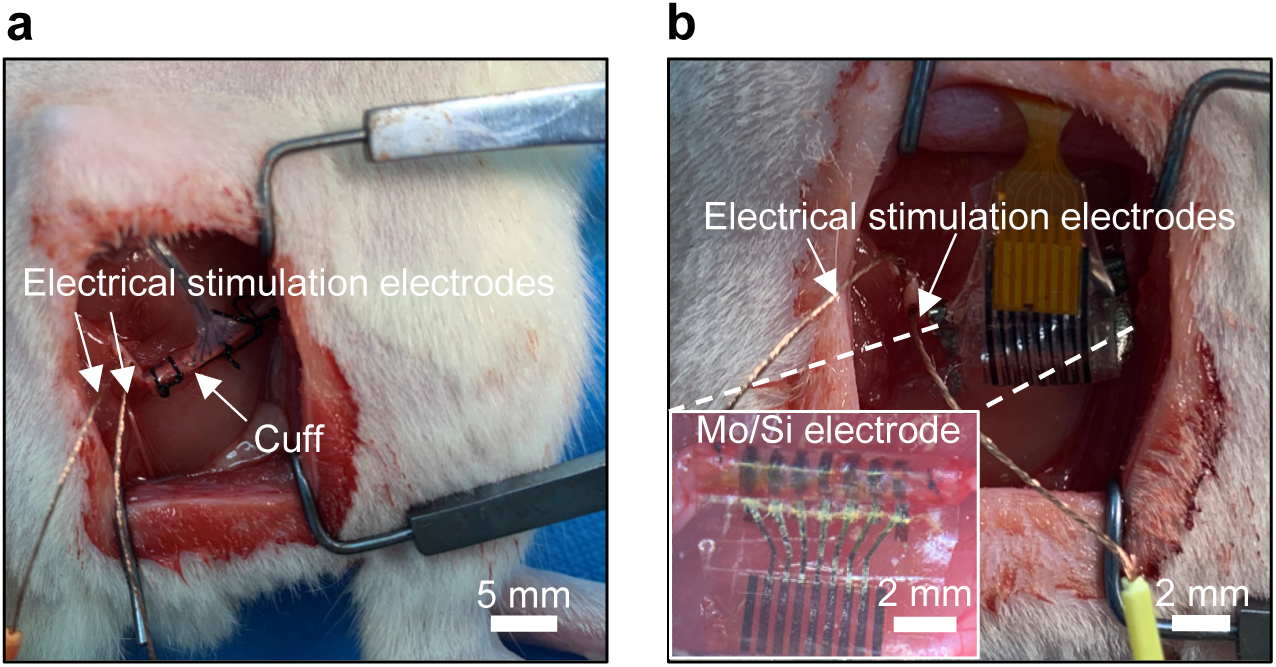
Evoked neural signals recorded by a, cuff electrode and b, biodegradable Mo/Si electrode array.

**Figure S6.**
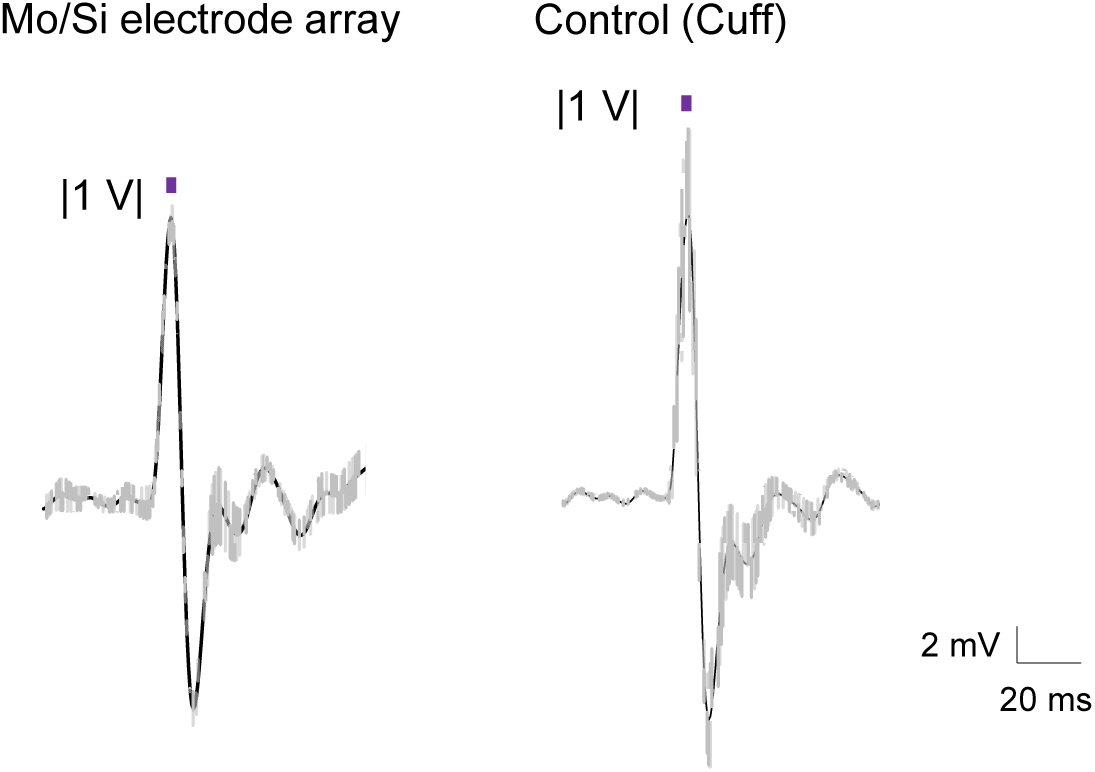
Evoked neural signals recorded by the Mo/Si electrode array and a commercial cuff electrode as a control. Stimulation frequency: 1 Hz.

**Figure S7.**
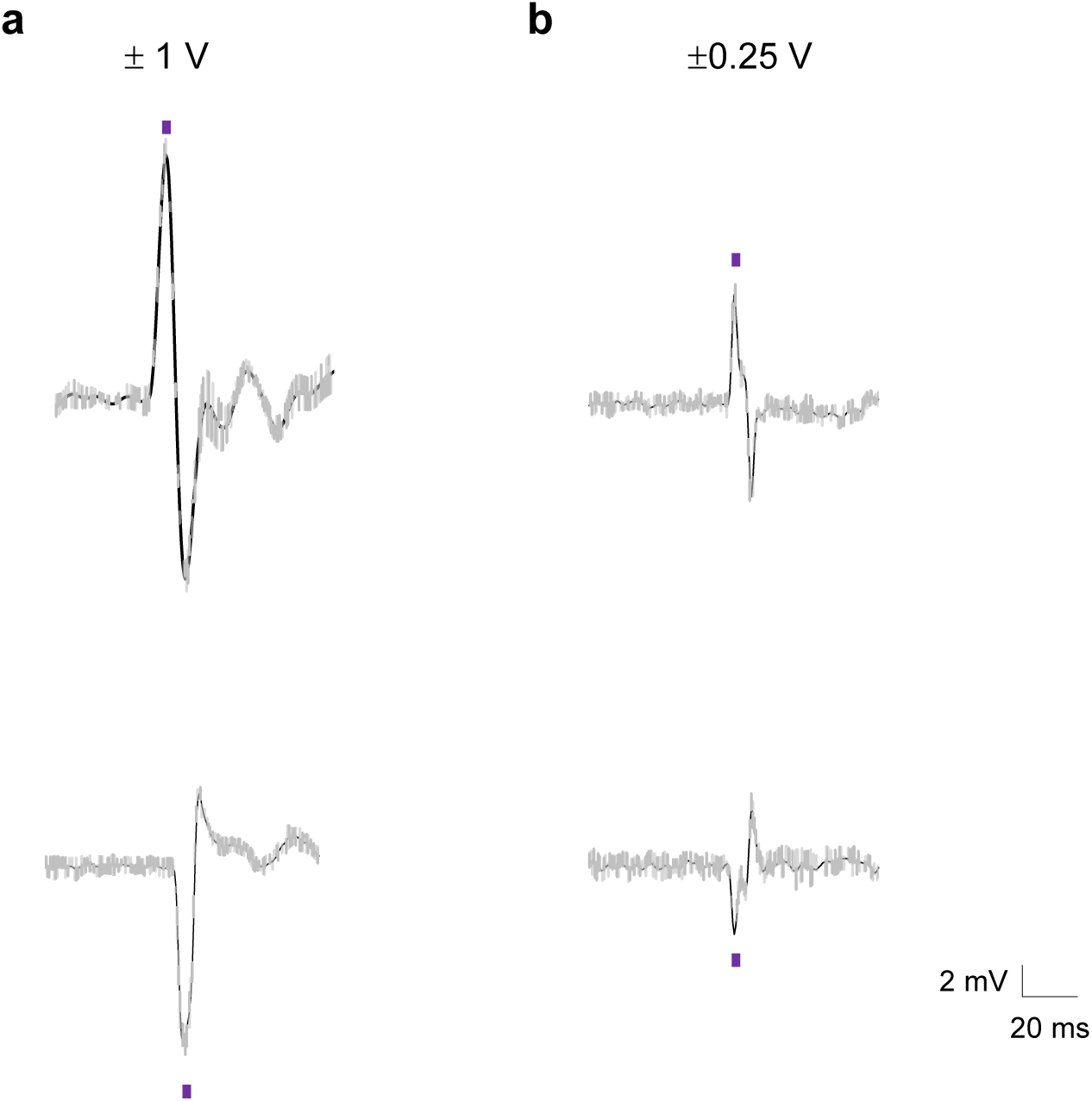
Evoked neural signals recorded by the Mo/Si electrode array with **a**, (1 V and **b**, (0.25 V. Stimulation frequency: 1 Hz.

**Figure S8.**
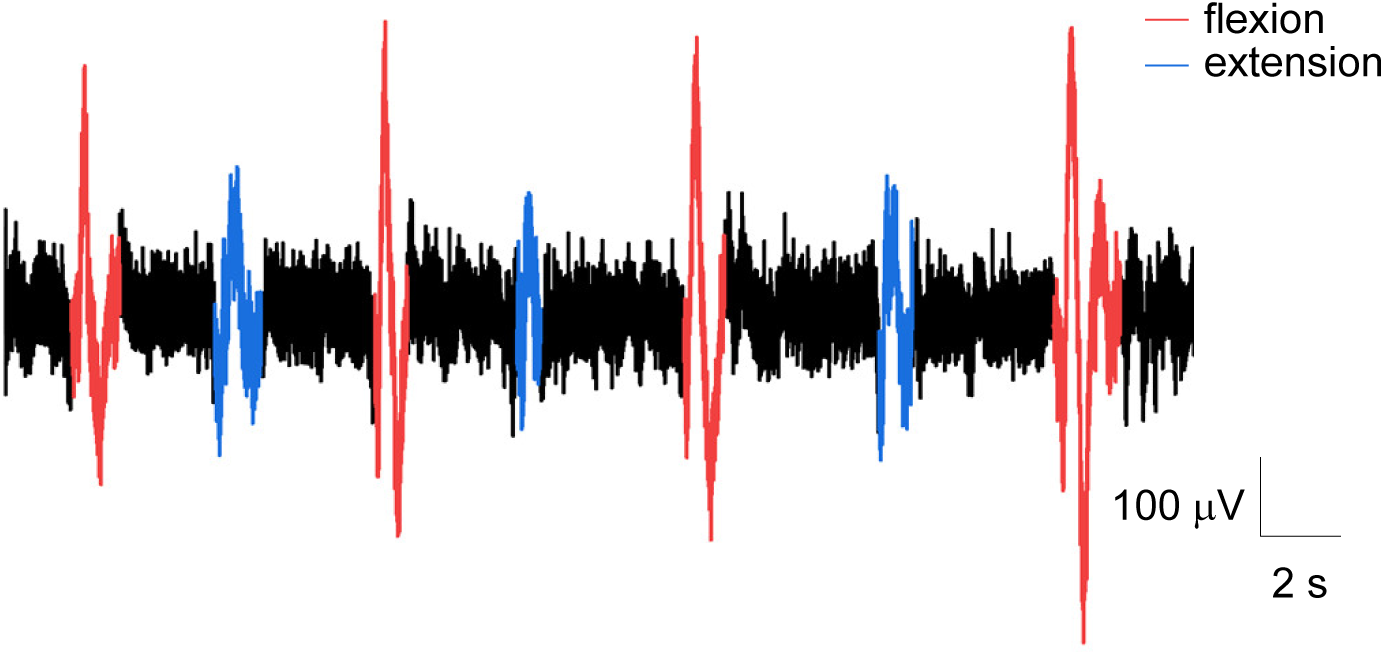
Recorded signal traces using cuff electrodes on sciatic nerves during the flexion-extension movement of the hindlimb.

**Figure S9.**
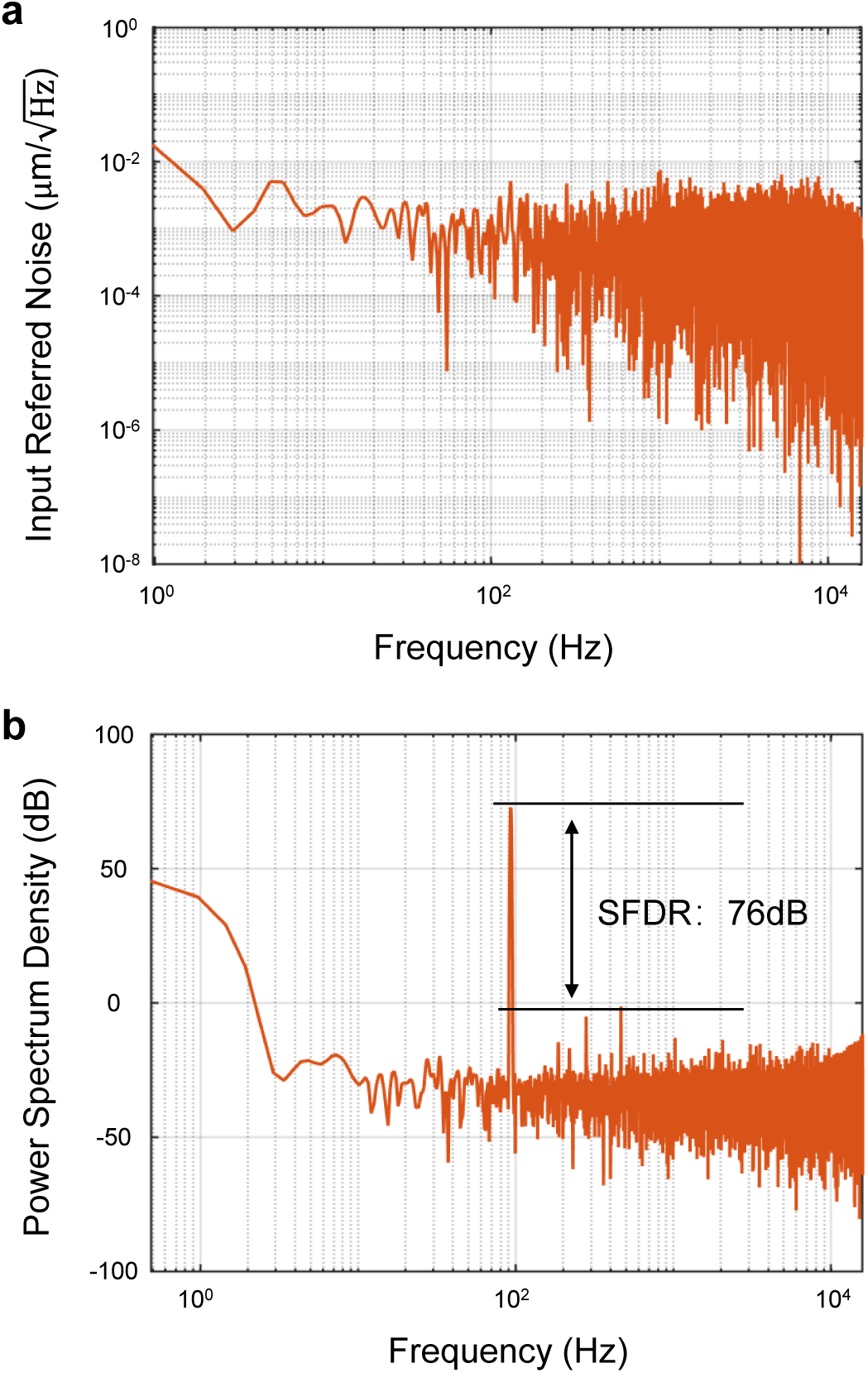
a, The measured input-referred-noise of the AFE chip. b, The measured SFDR of the AFE chip, with a 93Hz, 14 mVpp input signal.

**Figure S10.**
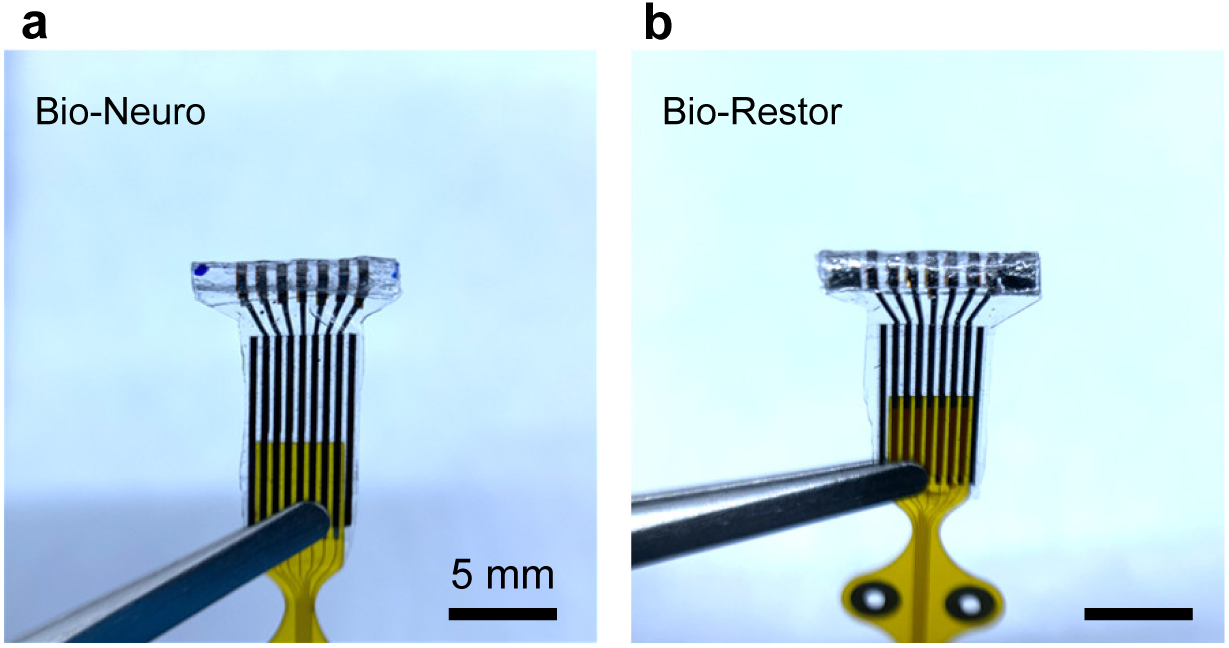
Images of a, Bio-Neuro (without the galvanic cell) and b, Bio-Restor.

**Figure S11.**
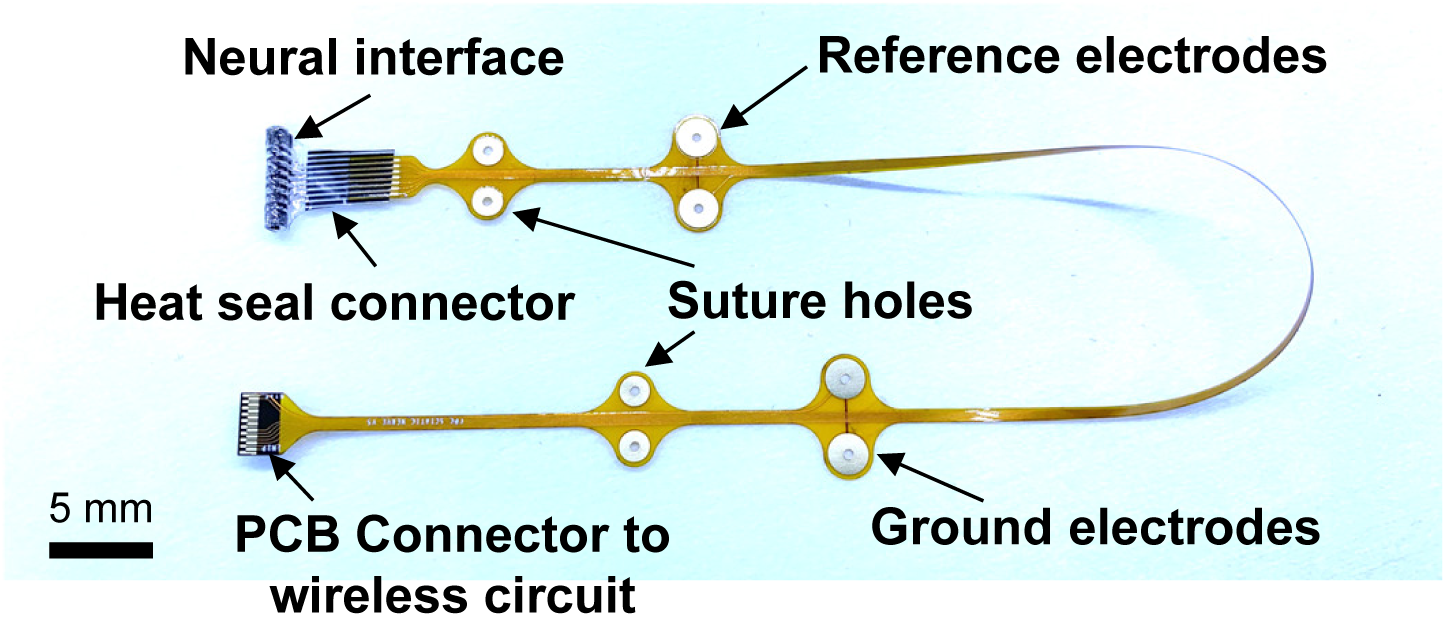
Structure of the flexible FPC connecting the neural interface and wireless circuit.

**Figure S12.**
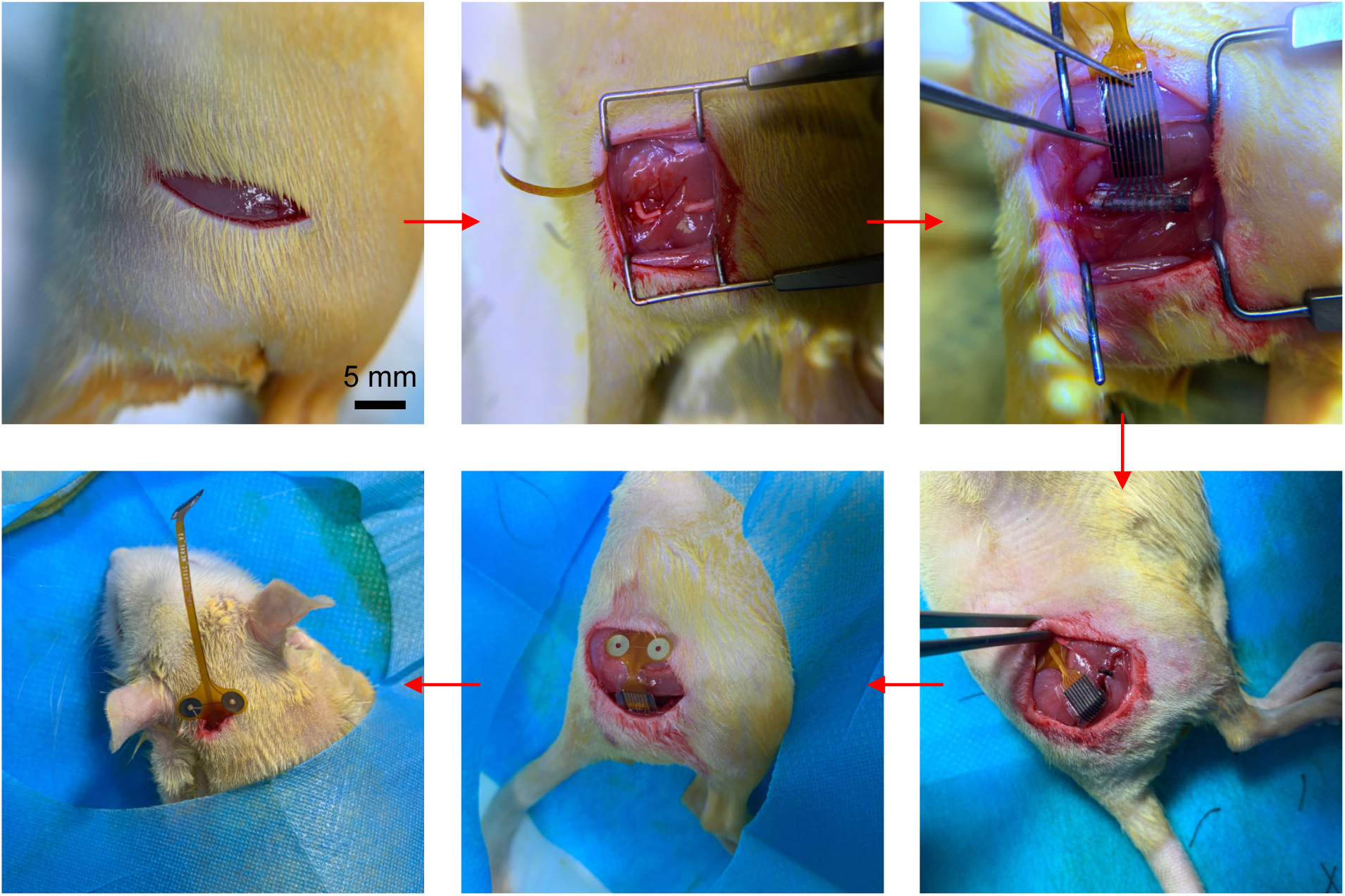
Implantation process of neural interfaces on the sciatic nerve in SD rats.

**Figure S13.**
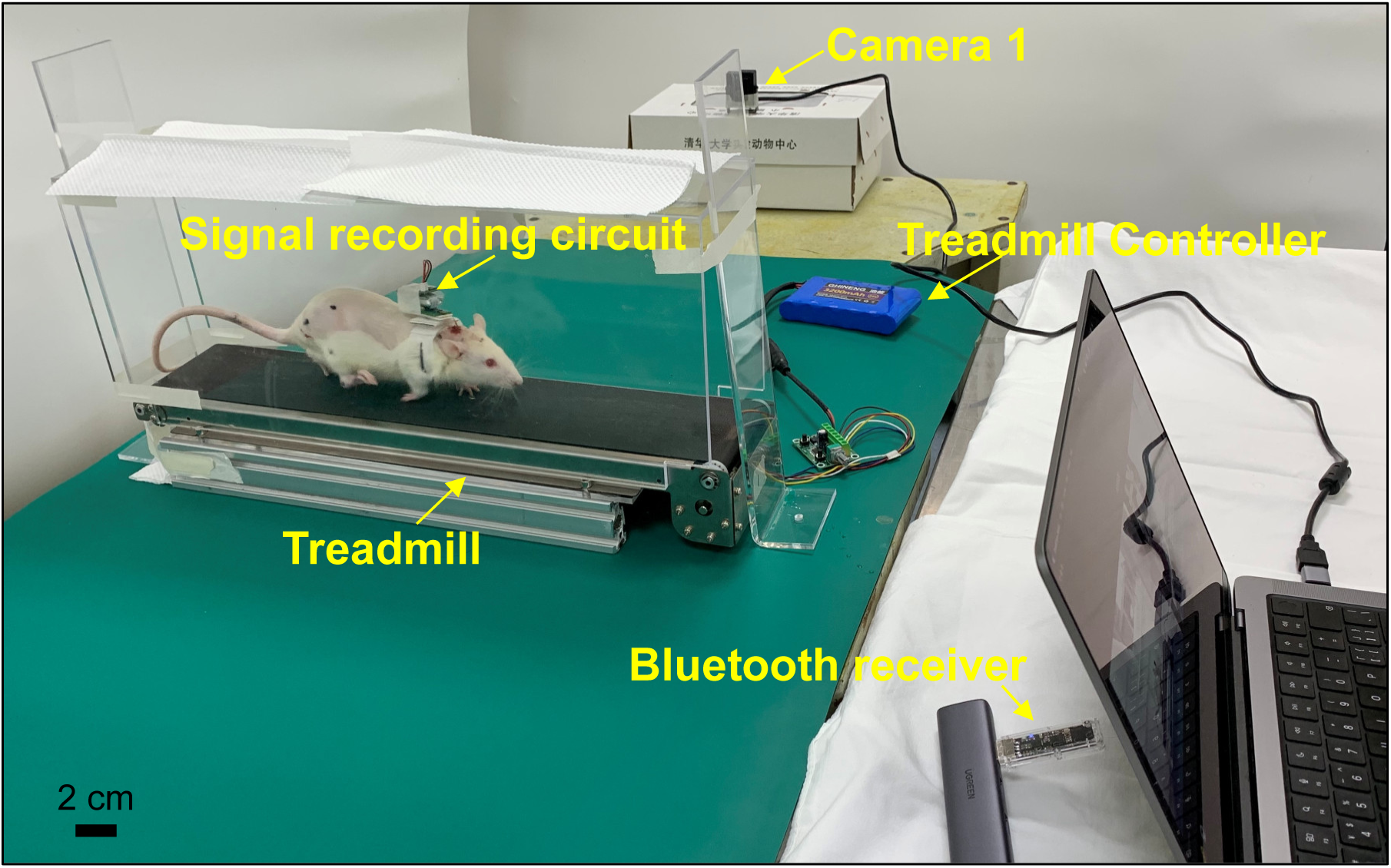
Setup for sciatic nerve signal recording and simultaneous gait analysis from lateral directions.

**Figure S14.**
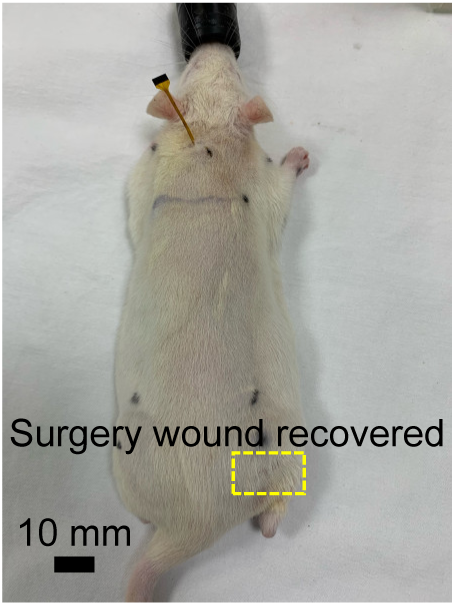
Image of a rat after 3 weeks implantation.

**Figure S15.**
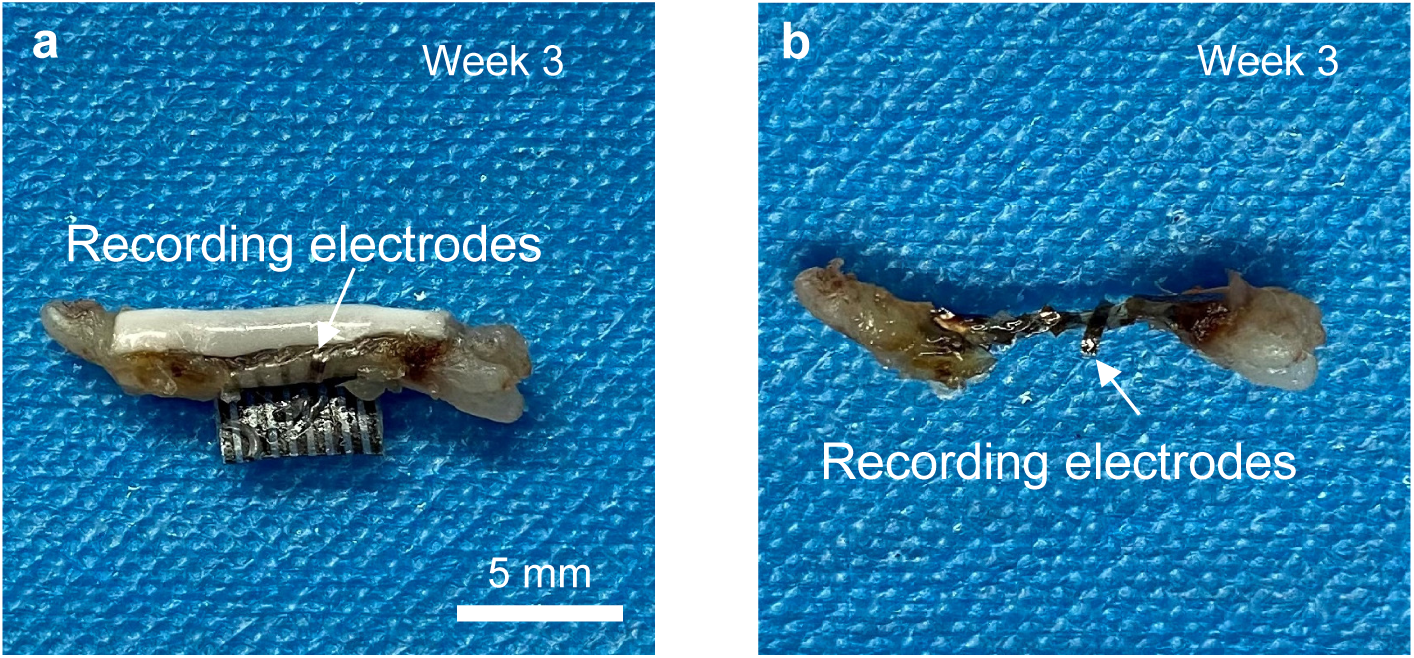
**a**, Intact Bio-Restor with regenerated nerve and **b**, the regenerated nerve has grown surrounding the recording electrodes at week 3.

**Figure S16.**
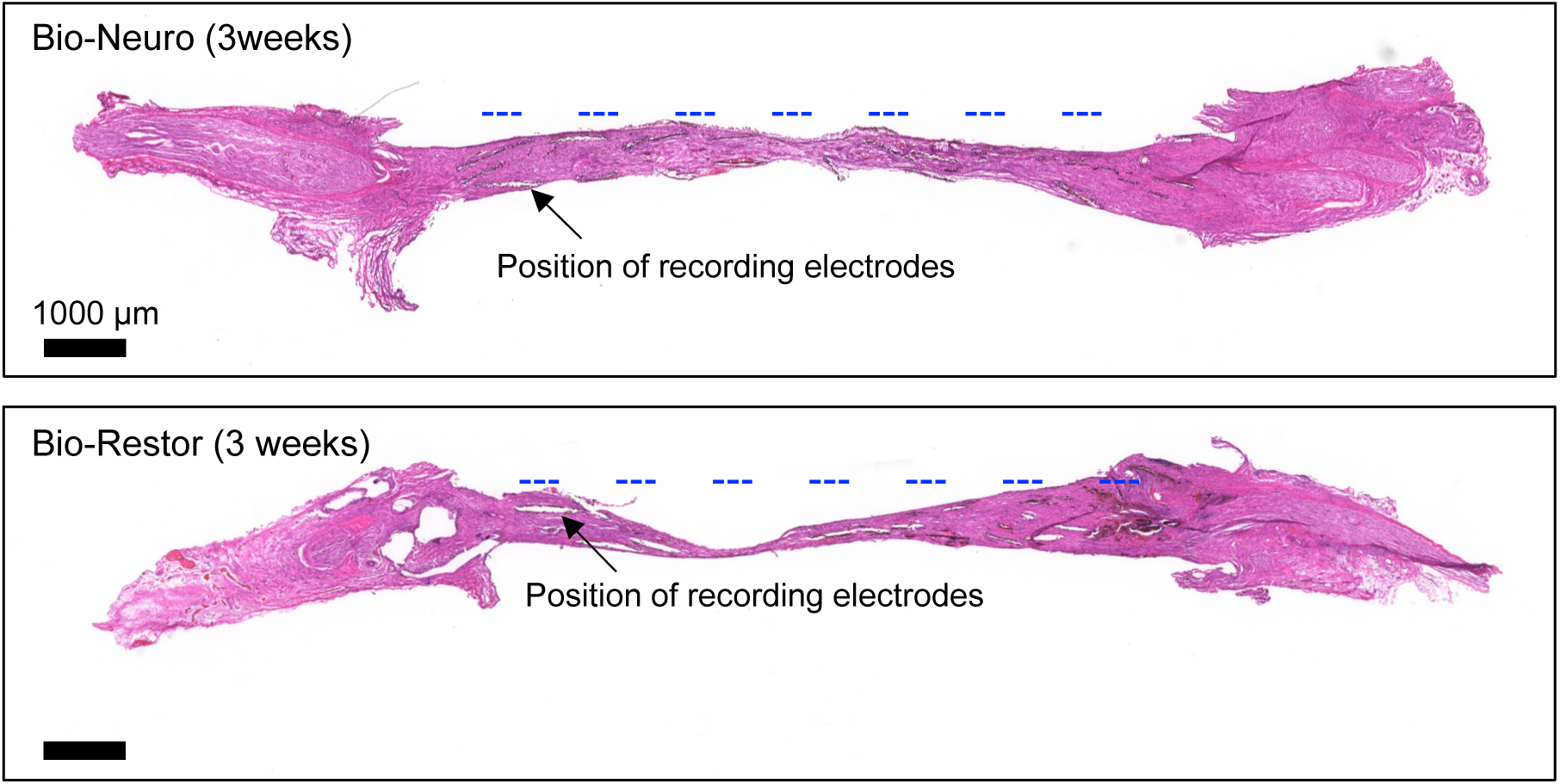
H&E staining images of longitudinal section of nerve tissues in the Bio-Restor and Bio-Neuro group at 3 weeks postimplantation. Blue dashed lines indicate the predicted position of recording electrodes.

**Figure S17.**
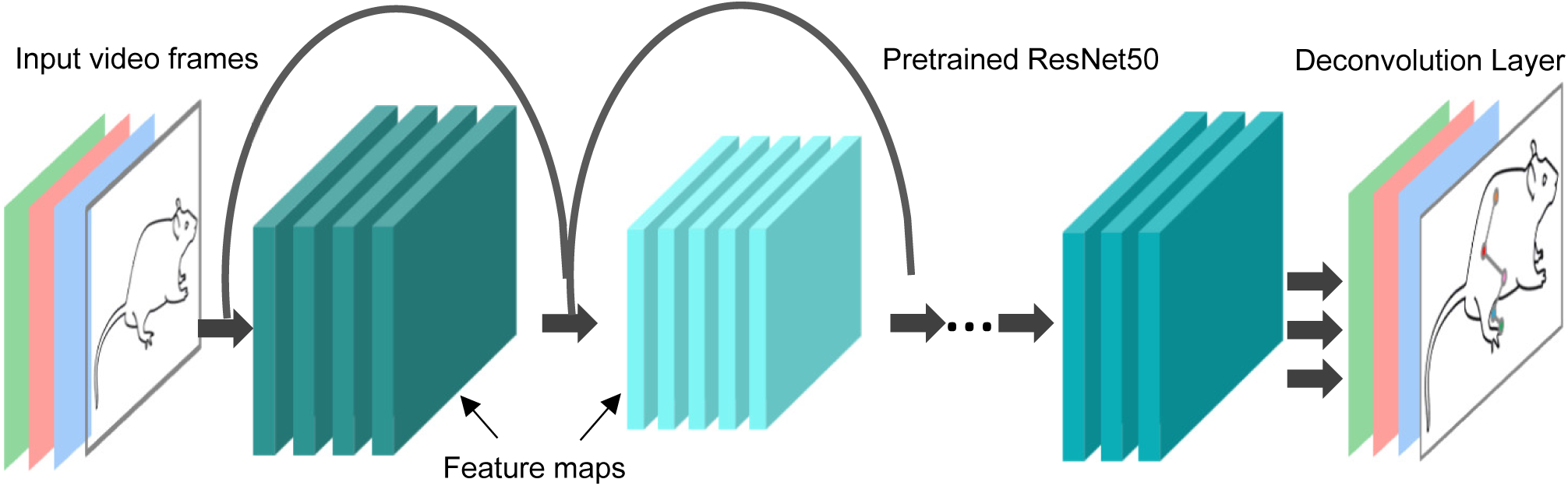
A deep neural architecture (DeepLabcut) to predict the location of body parts based on corresponding images with markers on the hip, knee, ankle, and toe.

**Figure S18.**
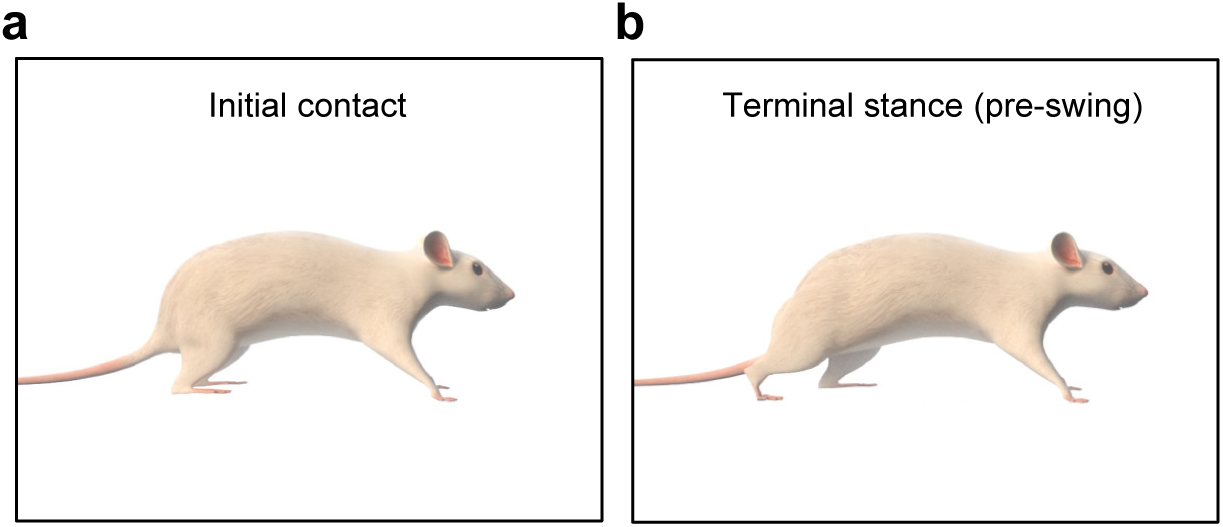
Image of the a, initial and b, terminal of stance phase (pre-swing).

**Figure S19.**
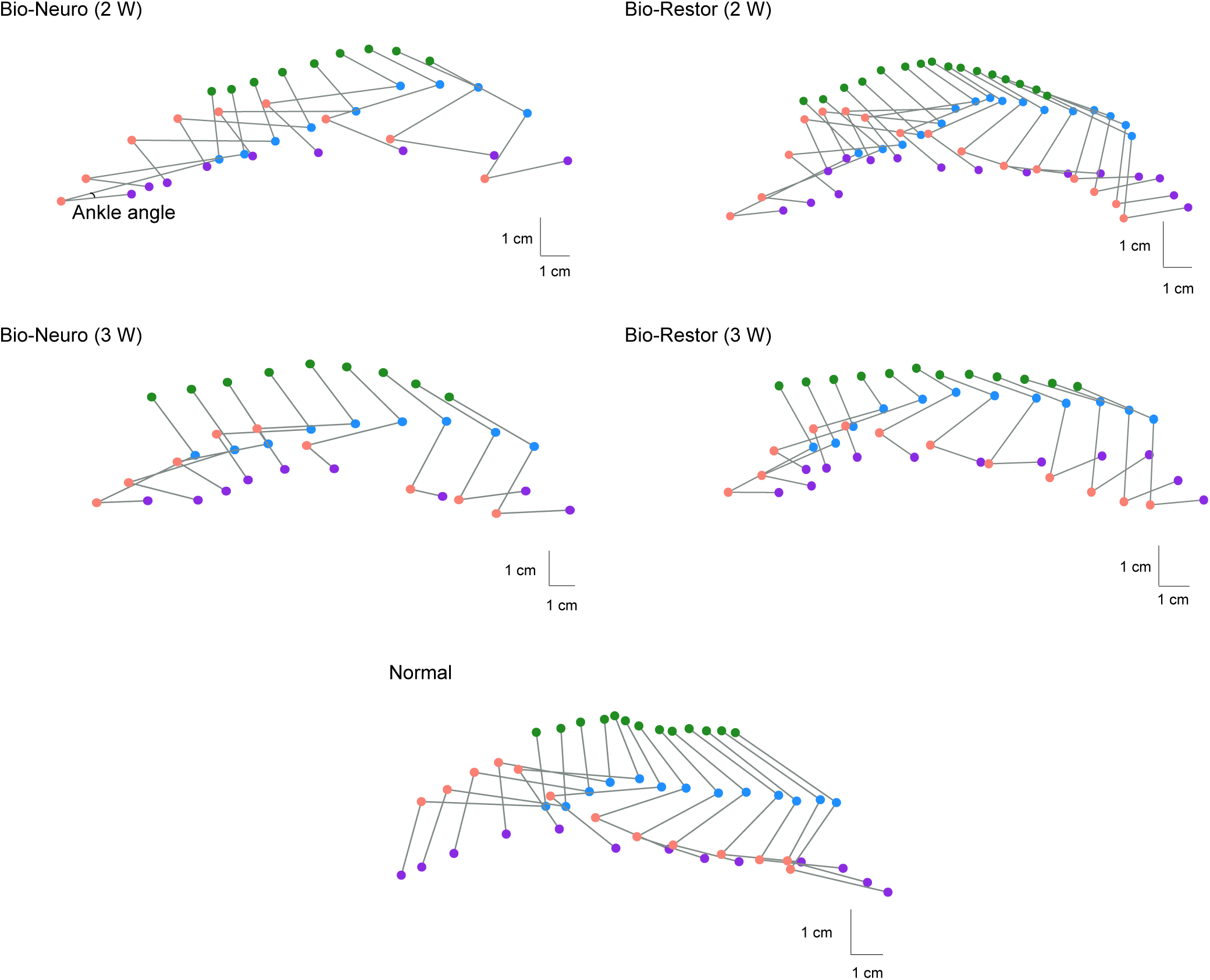
Stick diagram decompositions of hindlimb movements in Bio-Restor group, Bio-Neuro (2 and 3 weeks) after implantation and normal group.

**Figure S20.**
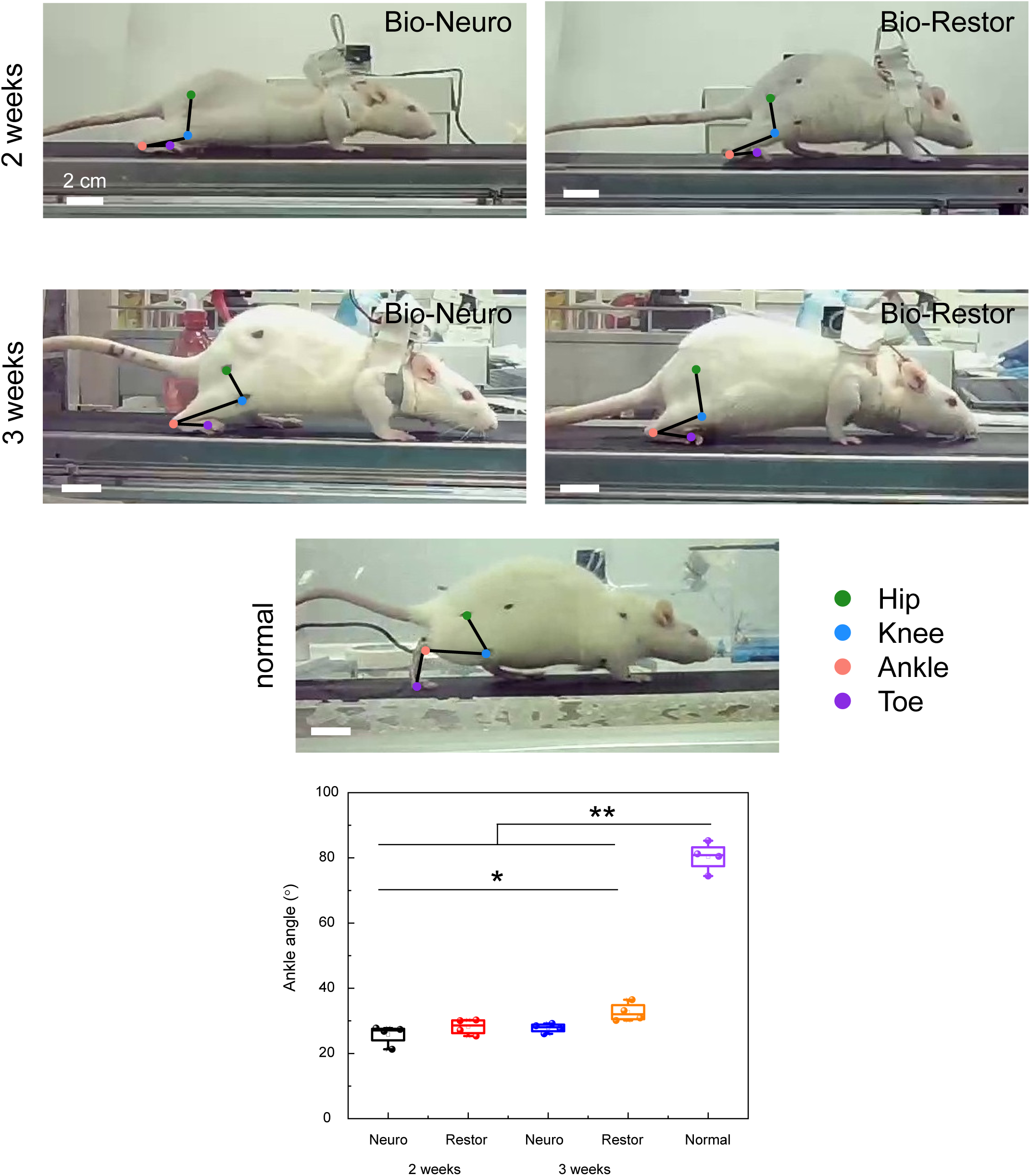
Representative images and statistics of ankle angle for Bio-Neuro, Bio-Restor and normal groups on terminal stance at 2 and 3 weeks after implantation.

**Figure S21.**
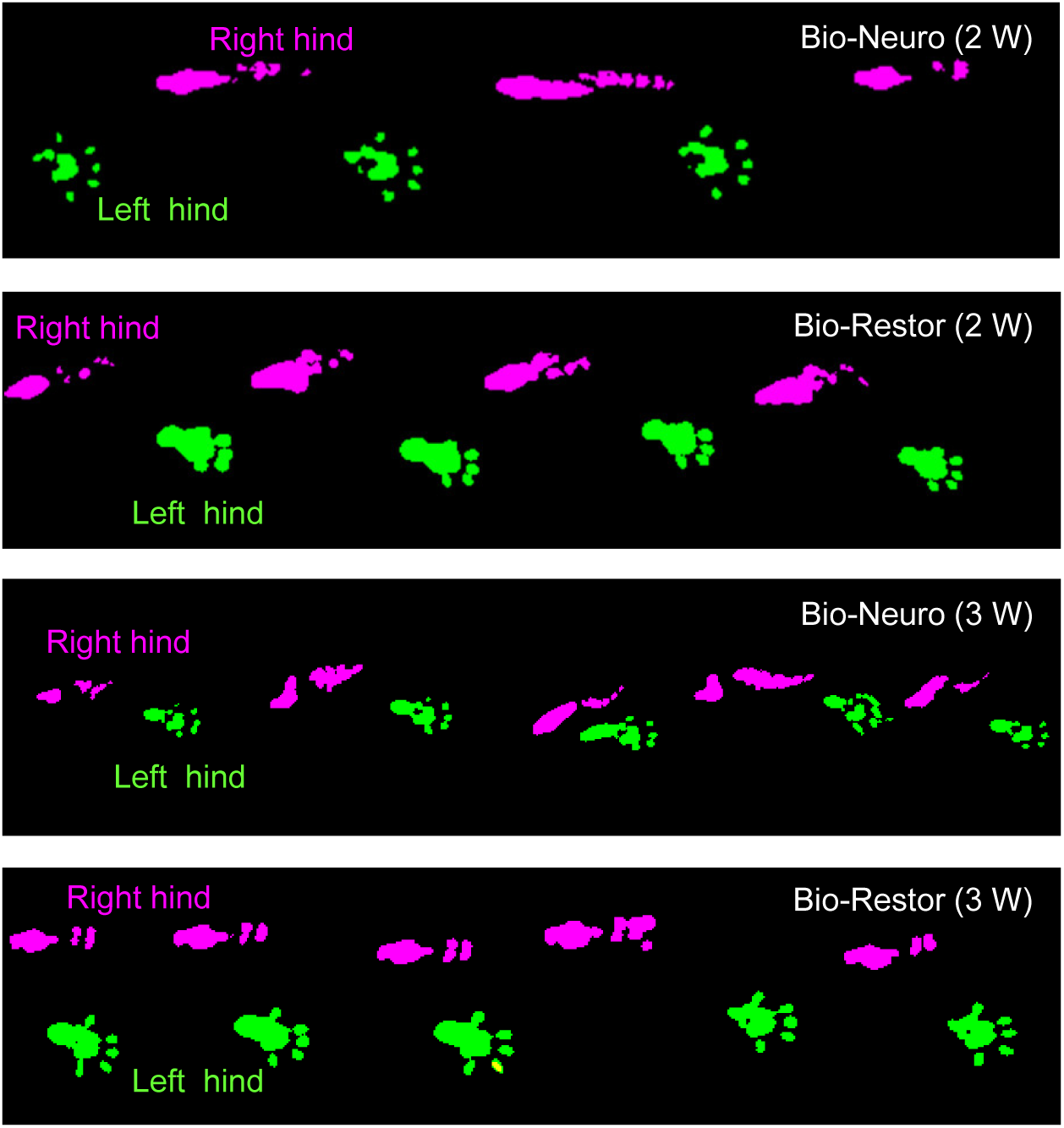
Representative hindlimb footprints for both injured (right hind) and contralateral sides (left hind) at 2 and 3 weeks after implantation.

**Figure S22.**
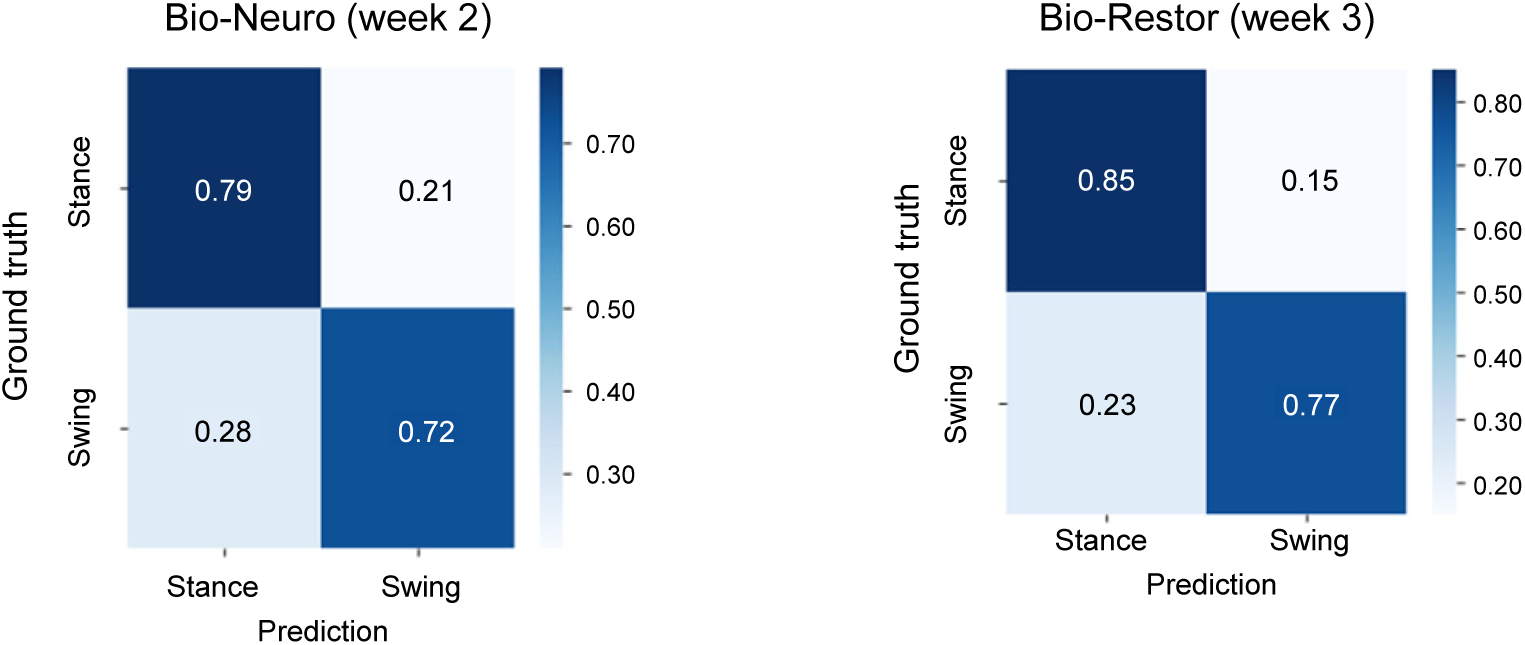
Confusion matrix of prediction and ground truth of swing and stance phase of the Bio-Neuro group at week 2 and Bio-Restor at week 3 postimplantation.

**Movie S1** The deployment of the device utilizing SMP on the intact nerve of a SD rat by applying PBS at 37 °C.

**Movie S2** Rats moving freely after device implantation.

**Movie S3** Representative recorded signal traces of the Bio-Neuro group at week 2. A correlation is observed between the patterns of electrical signals and the phases of walking or resting.

## Notes

### Competing Interest Statement

The authors have declared no competing interest.

